# Voltage-clamp fluorometry analysis of structural rearrangements of ATP-gated channel P2X2 upon hyperpolarization

**DOI:** 10.1101/2020.12.20.423705

**Authors:** Rizki Tsari Andriani, Yoshihiro Kubo

**Affiliations:** Division of Biophysics and Neurobiology, National Institute for Physiological Sciences, Aichi, Japan; Department of Physiological Sciences, The Graduate University for Advanced Studies, School of Life Science, Kanagawa, Japan

## Abstract

The gating of the ATP-activated channel P2X2 has been shown to be dependent not only on [ATP] but also on membrane voltage, despite the absence of a canonical voltage-sensor domain. We aimed to investigate the structural rearrangements of the rat P2X2 during ATP- and voltage-dependent gating by voltage-clamp fluorometry technique. We observed fast and linearly voltage-dependent fluorescence intensity (F) changes at Ala337 and Ile341 in the TM2 domain, which could be due to the electrochromic effect, reflecting the presence of a converged electric field here. We also observed slow and voltage-dependent F changes at Ala337, which reflect the structural rearrangements. Furthermore, we identified that the interaction between Ala337 in TM2 and Phe44 in TM1, located in close proximity in the ATP-bound open state, is critical for activation. Taken together, we propose that the voltage dependence of the interaction in the converged electric field underlies the voltage-dependent gating.

## Introduction

P2X2 is a member of the P2X receptor family, a ligand-gated cation channel which opens upon the binding of extracellular ATP (Brake et al., 1994; Valera et al., 1994). P2X receptors consist of 7 sub-classes (P2X1 – P2X7), in each of which subunits assemble to form trimeric homomers or heteromers (e.g. P2X2/P2X3) (Radford et al., 1997; North, 2002; L.-H. Jiang et al., 2003). Based on the solved crystal structures, P2X receptors are known to have a topology with two transmembrane (TM) domains (TM1 and TM2), a large extracellular ligand binding loop (ECD) where the ATP binding site is located, and intracellular N- and C-termini (Kawate et al., 2009; Hattori & Gouaux, 2012; Mansoor et al., 2016; McCarthy et al., 2019).

P2X2 is mainly distributed in smooth muscles, central nervous system (CNS), retina, chromaffin cells, and autonomic and sensory ganglia (Burnstock, 2003). Recent studies showed that P2X2 receptor expressed in hair cells and supporting cells has important roles in auditory transduction. A dominant negative polymorphism in human results in progressive hearing loss (Yan et al., 2013). Moreover, P2X2 in the cochlea is found to be involved in adaptation to elevated sound levels (Housley et al., 2013).

The P2X2 receptor has complex gating properties that consist of (1) the [ATP]-dependent gating, as well as (2) the voltage-dependent gating, in spite of the absence of a canonical voltage sensor domain, in clear contrast to typical voltage-gated ion channels, which have a voltage sensor domain (VSD) within their respective structures. In the presence of ATP, there is a gradual increase in the inward current upon hyperpolarization. The conductance – voltage relationship shifts toward depolarized potentials with an increase in [ATP]. Thus, the activation of the P2X2 channel is voltage-dependent as well as [ATP]-dependent (Nakazawa et al., 1997; Zhou & Hume, 1998; Nakazawa & Ohno, 2005; Fujiwara et al., 2009; Keceli & Kubo, 2009). Previous studies reported that this activation upon hyperpolarization is indeed an intrinsic property of the channel (Nakazawa et al., 1997; Zhou & Hume, 1998; Fujiwara et al., 2009).

It is of interest to know why and how P2X2 has voltage-dependent gating despite the absence of a canonical VSD. Previous studies extensively investigated the roles of amino acid residues in TM1 and TM2 during ATP-dependent gating and permeation (Haines et al., 2001; Jiang et al., 2001; Li et al., 2004; Khakh & Egan, 2005; Cao et al., 2007; Samways et al., 2008; Cao et al., 2009). In contrast, information about amino acid residues, particularly in TM domains, which might play important roles during voltage-dependent gating is still limited. A previous study identified positively-charged amino acid residues in the ATP binding pocket (K69, K71, R290, and K308; *r*P2X2 numbering) and aromatic amino acid residues in TM1 (Y43, F44, and Y47; *r*P2X2 numbering) which are critical for ATP- and voltage-dependent gating of P2X2 receptor (Keceli & Kubo, 2009). However, those residues were not the sole determinant of [ATP]- and voltage-dependent gating of the P2X2 receptor. The interpretation as to the mechanism is not yet straightforward, and thus, the key amino acid residue that has a major contribution to the voltage sensing mechanism in P2X2 receptor is yet to be discovered.

Moreover, the details of the structural rearrangements upon ATP binding in the pore region remain controversial, due to discrepancies between the *zf*P2X4 structural data and P2X experimental data (Kawate et al., 2009; Kracun et al., 2010; Li et al., 2010; Hattori & Gouaux, 2012; Heymann et al., 2013; Habermacher et al., 2016), as well as between the solved crystal structures of TM domains of *zf*P2X4 and *h*P2X3. The comparison highlights longer TM domains and visualized cytoplasmic domain in *h*P2X3 (Kawate et al., 2009; Hattori & Gouaux, 2012; Mansoor et al., 2016). The structural study of *h*P2X3 visualized a region called the cytoplasmic cap in the ATP-bound open state, and it was further confirmed by the *r*P2X7 crystal structure (Mansoor et al., 2016; McCarthy et al., 2019). Thus, the present study aims at analyzing the structural rearrangements of the P2X2 receptor upon (1) ATP- and (2) voltage-dependent gating, by voltage-clamp fluorometry (VCF) using a fluorescent unnatural amino acid (fUAA) as a probe.

The combination of fluorometry and voltage-clamp recording offers a powerful method to track down real time conformational changes within the ion channel structure (Mannuzzu et al., 1996; Cha & Bezanilla, 1997; Pless & Lynch, 2008; Nakajo & Kubo, 2014; Talwar & Lynch, 2015). The use of fUAA as a probe made it possible to label any residues within the protein, including those at the lower TM and intracellular regions, which will not be accessible by conventional VCF fluorophores such as Alexa-488 maleimide (Kalstrup & Blunck, 2013; Sakata et al., 2016; Kalstrup & Blunck, 2018; Klippenstein et al., 2018). Moreover, a direct incorporation of the fUAA will increase the labelling efficiency and also prevent non-specific labelling (Kalstrup & Blunck, 2013; Sakata et al., 2016).

The fUAA used here, *3-(6-acetylnaphthalen-2-ylamino)-2-aminopropionic acid* (Anap), was incorporated into the *r*P2X2 protein by using a non-sense suppression method where the tRNA Anap-CUA and tRNA-synthetase pair is used to introduce Anap at an amber nonsense codon mutation (Lee et al., 2009; Chatterjee et al., 2013; Klippenstein et al., 2018), as shown in **Fig. 1A**. By performing VCF recording using Anap as a fluorophore, we analyzed the structural dynamics of the P2X2 receptor undergoing complex gating. In the present study, we observed evidence of voltage-dependent conformational changes around the transmembrane regions. We also investigated the key amino acid residues in each TM region whose interaction might have major contributions to the ATP- and voltage-dependent gating of the P2X2 receptor.

**Figure 1.**
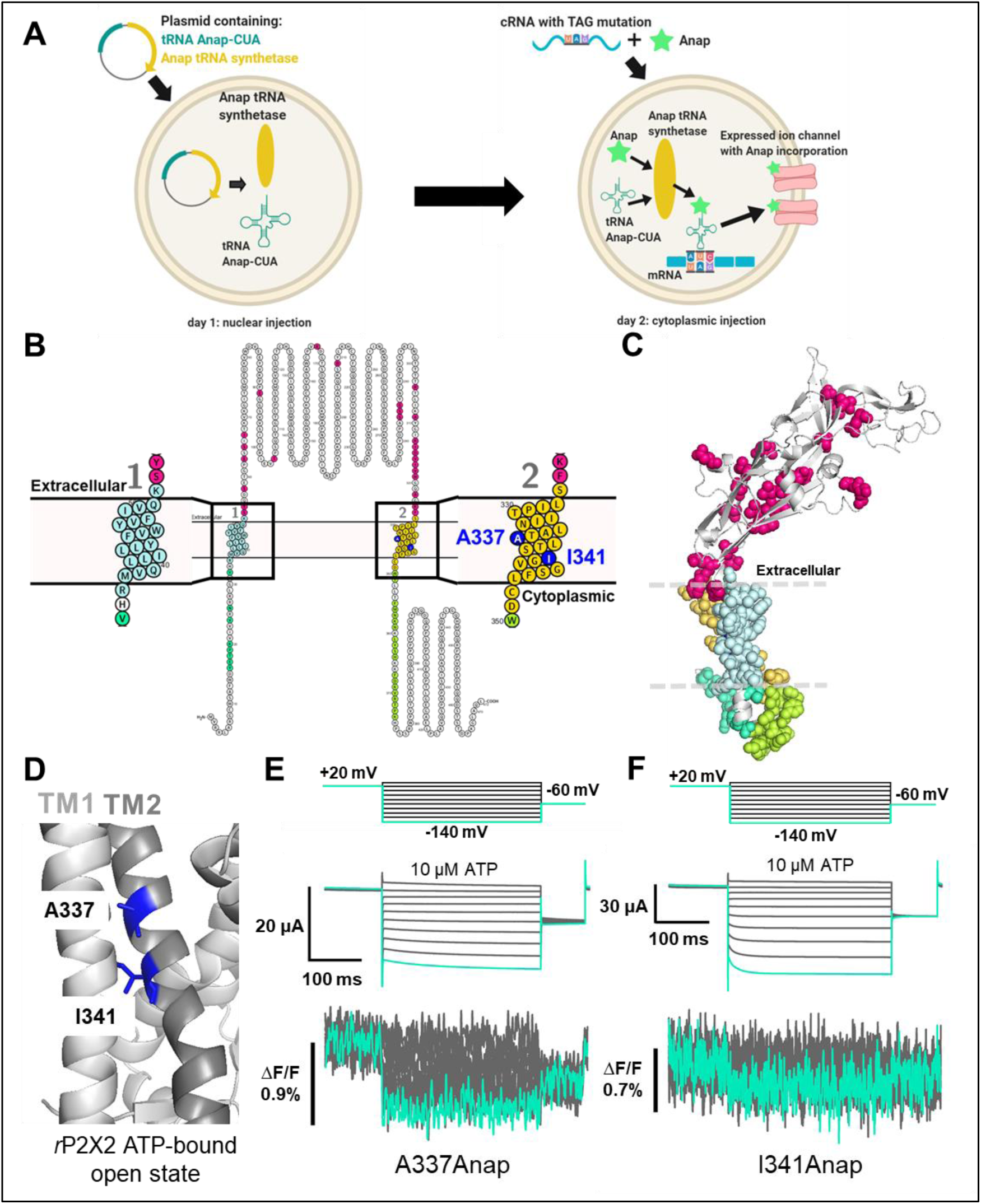
Fluorescence signal changes of Anap incorporated P2X2 receptor evoked by ATP and voltage. (**A**) A scheme depicting the principle of the direct incorporation of fUAA (Anap) into the ion channel protein. The plasmid containing tRNA Anap-CUA and tRNA synthase is injected into the nucleus of *Xenopus laevis* oocytes. On the following day, channel cRNA with TAG mutation is co-injected with Anap into the cytoplasmic region of the oocytes. Anap-incorporated channel protein was expressed successfully after the optimum incubation period. (**B, C**) A scheme to visualize the Anap scanning regions by individual amino acid residue representation (**B**) and within the protein structure (**C**), respectively. Anap mutant scanning was done by introducing TAG mutation one at a time in all regions of the P2X2 receptor (a total of 96 positions), which include the N-terminus (8 positions, turquoise), TM2 (24 positions, yellow), extracellular domain (ECD, 25 positions, magenta), TM1 (20 positions, light blue), and C-terminus (19 positions, lime green). Voltage-dependent fluorescence changes of Anap were observed only at A337 and I341 in the TM2 domain (colored by dark blue). (**D**) The sites of the introduced TAG mutations, A337 and I341 in the TM2 domain, which gave voltage-evoked fluorescence changes. All of *r*P2X2 structure representations in (**C**) and (**D**) were based on homology modeling from the ATP-bound open state *h*P2X3 crystal structure data (PDB ID: 5SVK; Mansoor et al., 2016). (**E, F**) Representative current traces and fluorescence signal upon ATP and voltage application in Anap mutants (A337: ΔF/F=0.5±0.2% at 440 nm, n=3; I341: ΔF/F=0.3±0.2% at 440 nm, n=3; respectively). (**Source Data 1 Figure 1**)

## Results

### Fluorescence signal changes of Anap-labeled P2X2 receptor evoked by ATP and voltage

As the P2X2 receptor does not have a canonical voltage-sensing domain (VSD), we performed Anap scanning by introducing TAG mutations one at a time in all regions of the P2X2 receptor, including the cytoplasmic N-terminus (8 positions), TM1 (20 positions), ECD, where the ATP binding site is located (25 positions), TM2 (24 positions), and cytoplasmic C-terminus (19 positions) (**Fig. 1B, C**). The whole of TM1 and TM2 was scanned, as these are the transmembrane domains in which a non-canonical voltage sensor might be located.

From the total of 96 positions of Anap mutant scanning in the P2X2 receptor, many showed ATP-evoked fluorescence intensity changes (**Supplementary Table 1**). As major and overall structural movement occurs upon the binding of ATP during the channel’s transition from closed to open state in the P2X receptor (Kawate et al., 2009; Hattori & Gouaux, 2012; Mansoor et al., 2016; McCarthy et al., 2019), the results go well with the expectation that ATP-evoked fluorescence change would be observed at many positions labeled by Anap.

In contrast, at only two positions located at TM2 domain, out of 96 scanning positions, could we detect Anap fluorescence intensity changes (ΔF) in response to voltage stimuli. The two positions are A337 (ΔF/F=0.5±0.2% upon voltage change from +40 mV to −140 mV at 440 nm, n=3, **Fig. 1D, E**) and I341 (ΔF/F=0.3±0.2% upon voltage change from +40 mV to −140 mV at 440 nm, n=3, **Fig. 1D, F**). Although the Anap ΔF were observed after the application of 10 μM ATP and voltage step pulses, there are two major concerns as follows: (1) ΔF is close to the limit of detection because signal to noise ratio is low, making it hard to perform further analysis e.g. F-V relationship; (2) The incidence of fluorescence change detection is also low. Thus, at this point, further analysis to determine the structural rearrangements with which Anap ΔF is associated could not be performed.

### SIK inhibitor treatment improved VCF optical signal in Anap labeled *Ci*-VSP and P2X2 receptor

To overcome the problems of only small fluorescence changes and low incidence of successful detection of fluorescence changes, a small molecule kinase inhibitor, namely an SIK inhibitor (HG-9-91-01), was applied by injection into the oocytes, to decrease the intrinsic background fluorescence (Lee & Bezanilla, 2019). This inhibitor promotes UV-independent skin pigmentation, by increasing the production of melanin (Mujahid et al., 2017), resulting in a darker surface of the animal pole of the oocyte. As the intrinsic background fluorescence of the oocytes is decreased, the percentage of fluorescence change (ΔF/F) is expected to increase.

Optimization of SIK inhibitor treatment in VCF experiments using Anap as fluorophore was achieved for the following conditions. (1) The optimal concentration of SIK inhibitor injected into the oocyte to give the maximum effect of decreasing the intrinsic background fluorescence. (2) The optimal injection conditions for the location of the microinjection into the oocyte (nuclear or cytoplasmic) and the duration of incubation.

*Ci-*VSP F401Anap (Sakata et al., 2016), was used as a positive control to obtain reproducible and distinct results (**Fig. 2A – E**). Oocytes were pre-treated with two concentrations of SIK inhibitor (30 nM and 300 nM, reflecting the concentration of injected solution). 300 nM SIK application increased ΔF/F more than twice that of non-treated oocytes, whereas the application of 30 nM did not give a significant increase (ΔF/F= 10.6%±2.5 at 500 nm, n=6; ΔF/F= 3.2%±0.8, n=8; and ΔF/F= 6.4%±1.9, n=6; respectively, **Fig. 2A – D**). This showed that 300 nM SIK inhibitor injected into the oocytes could decrease the intrinsic background fluorescence of the oocytes, thus increasing ΔF/F.

**Figure 2.**
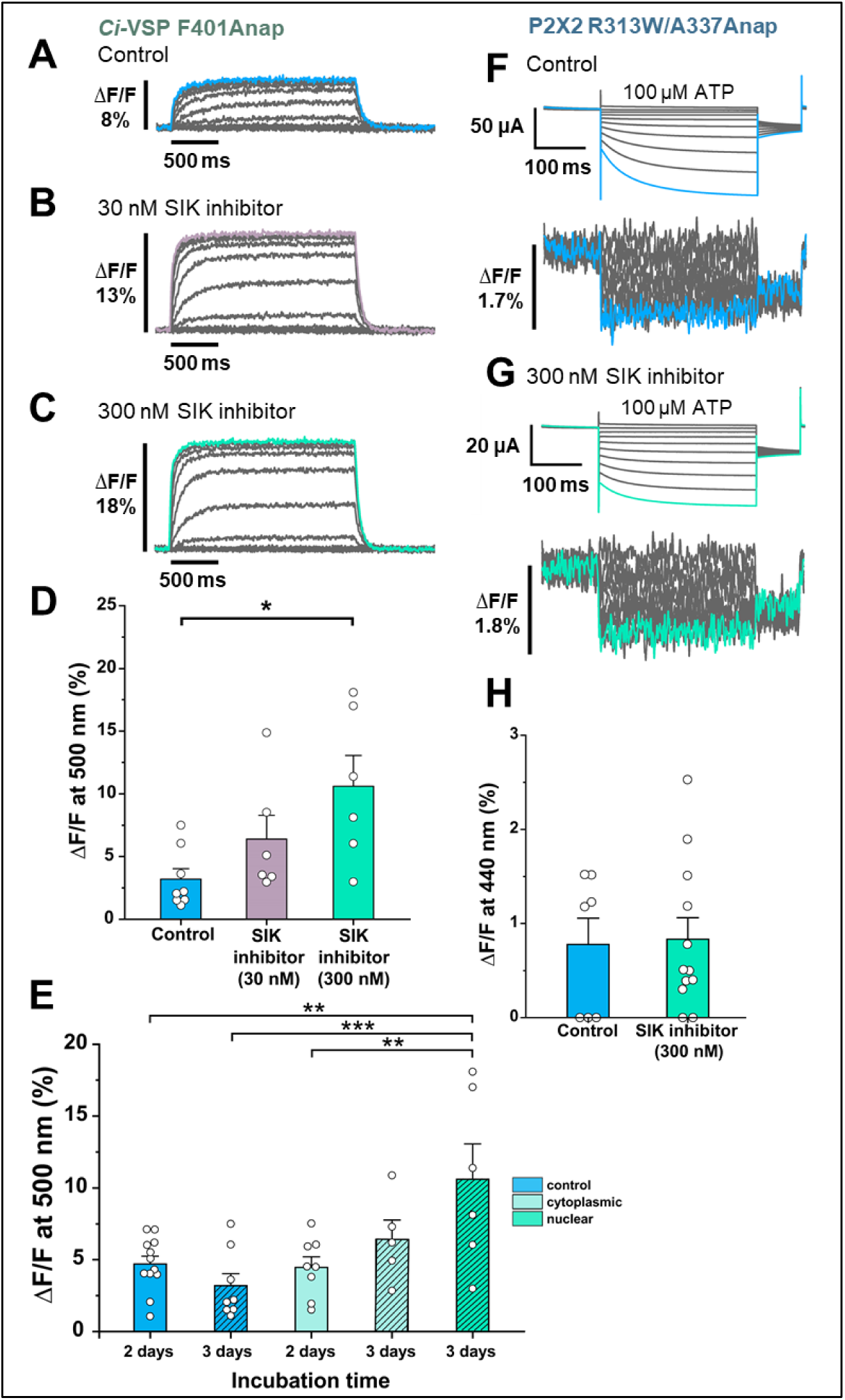
Effect of SIK inhibitor treatment in Anap incorporated *Ci*-VSP and P2X2 receptor. SIK inhibitor treatment improved the VCF optical signal. (**A - C**) Representative fluorescence signal of VCF recordings of *Ci*-VSP without SIK inhibitor treatment, with 30nM, and with 300 nM SIK inhibitor treatment (ΔF/F= 3.2%±0.8 at 500 nm, n=8; ΔF/F= 6.4%±1.9 at 500 nm, n=6; and ΔF/F= 10.6%±2.5 at 500 nm, n=6 respectively). (**D**) Comparison of non-treated (control) group (n=8), 30 nM (n=6), and 300 nM SIK inhibitor application (n=6); *, p≤0.05, p=0.016, one-way ANOVA with Tukey’s post-hoc test for 300 nM, compared to the control group. (**E**) Comparison of the incubation time and site of injection of SIK inhibitor treatment using 300 nM SIK inhibitor: control group, 2 days incubation (n=12), control group, 3 days incubation (n=8), SIK inhibitor treatment with cytoplasmic injection with 2 days incubation (n=8), with cytoplasmic injection for 3 days (n=5), with nuclear injection for 3 days (n=6); **, p≤0.01, ***, p≤0.001, one-way ANOVA with Tukey’s post-hoc test. (**F, G**) Representative current traces and fluorescence signal of VCF recordings of P2X2 receptor (A337Anap/R313W) without SIK inhibitor treatment and with the application of 300 nM SIK inhibitor (ΔF/F= 0.77%±0.3 at 440 nm, n=7; and ΔF/F= 0.83%±0.2 at 440 nm, n=12, respectively). (**H**) A comparison of non-treated (control) group (n=7) and 300 nM SIK inhibitor application (n=12) (p = 0.881, two sample t-test for 300 nM compared to the control group). All error bars are ± s.e.m centered on the mean. (**Source Data 1 Figure 2**)

Subsequently, a second series of optimization experiments was performed. In all of the following experiments, 300 nM SIK inhibitor was used. Control groups consisted of non-treated oocytes which were incubated for either 2 or 3 days, resulting in ΔF/F=4.7%±0.5 (n=12) and ΔF/F=3.2%±0.8 (n=8), respectively. The nuclear injection group, which was incubated for 3 days, had a larger ΔF/F than the other groups (ΔF/F= 10.6%±2.5 at 500 nm, n=6). The cytoplasmic injection groups, which were incubated for either 2 or 3 days, resulted in ΔF/F=4.5%±0.7 (n=8) and ΔF/F=6.4%±1.3 (n=5) respectively. These results suggest that the optimal conditions for SIK inhibitor treatment are nuclear injection with 300 nM SIK inhibitor and 3 days incubation (**Fig. 2E**).

After the optimal concentration, injection method, and incubation period were determined for the *Ci*-VSP experiment, the SIK inhibitor was then applied to the P2X2 A337Anap/R313W mutant (**Fig. 2F, G**). R313W is a mutation which decreases the basal current in the absence of ATP, and the details are described later in **Fig. 4 and Fig. 4—figure supplement 1**. 300 nM SIK inhibitor treatment did not make any significant difference, in terms of the percentage of the fluorescence change compared to the control group (ΔF/F= 0.77%±0.3 at 440 nm, n=7 and ΔF/F= 0.83%±0.2 at 440 nm, n=12, respectively, **Fig. 2H**). However, in the analysis of the incidence of detectable ΔF of Anap, the group treated with 300 nM SIK inhibitor showed a higher incidence than the control group (control = 57%, n=7; 300 nM SIK inhibitor application = 80%, n=12; **Fig. 2—figure supplement 1. A, B**). These results showed that in the case of P2X2, SIK inhibitor treatment improved the incidence of detectable ΔF/F. Therefore, we decided to use the SIK inhibitor in all of the following experiments.

### ATP- and voltage-evoked Anap fluorescence changes at A337 and I341 in TM2 exhibit a fast kinetics and linear voltage-dependence

By the application of 300 nM SIK inhibitor, a more frequent and improved signal-to-noise ratio of Anap ΔF could be observed at A337 (ΔF/F= 1.5%±0.2 at 440 nm, n=8, **Fig. 3A**). VCF recordings were performed by the application of 10 μM ATP and voltage step pulses from +40 mV to −140 mV with a holding potential at +20 mV. Fluorescence intensity change occurred almost instantaneously in less than 5 ms (**Fig. 3B**). This showed that the kinetics of ΔF/F are very rapid and faster than the time course of the voltage-dependent current activation. This also correlates well with the speed of the actual membrane potential change achieved by voltage clamp. Besides, the ΔF/F – V relationship of A337Anap showed a linear voltage-dependence (y = 0.011x + 0.016; R² = 0.99, n=8, **Fig. 3C**) in the recorded voltage range. These analyses of fluorescence changes at A337 indicated that the downward fluorescence change is not associated with the protein conformational change. These changes are rather thought to be well explained as a phenomenon related to electrochromic effect.

**Figure 3.**
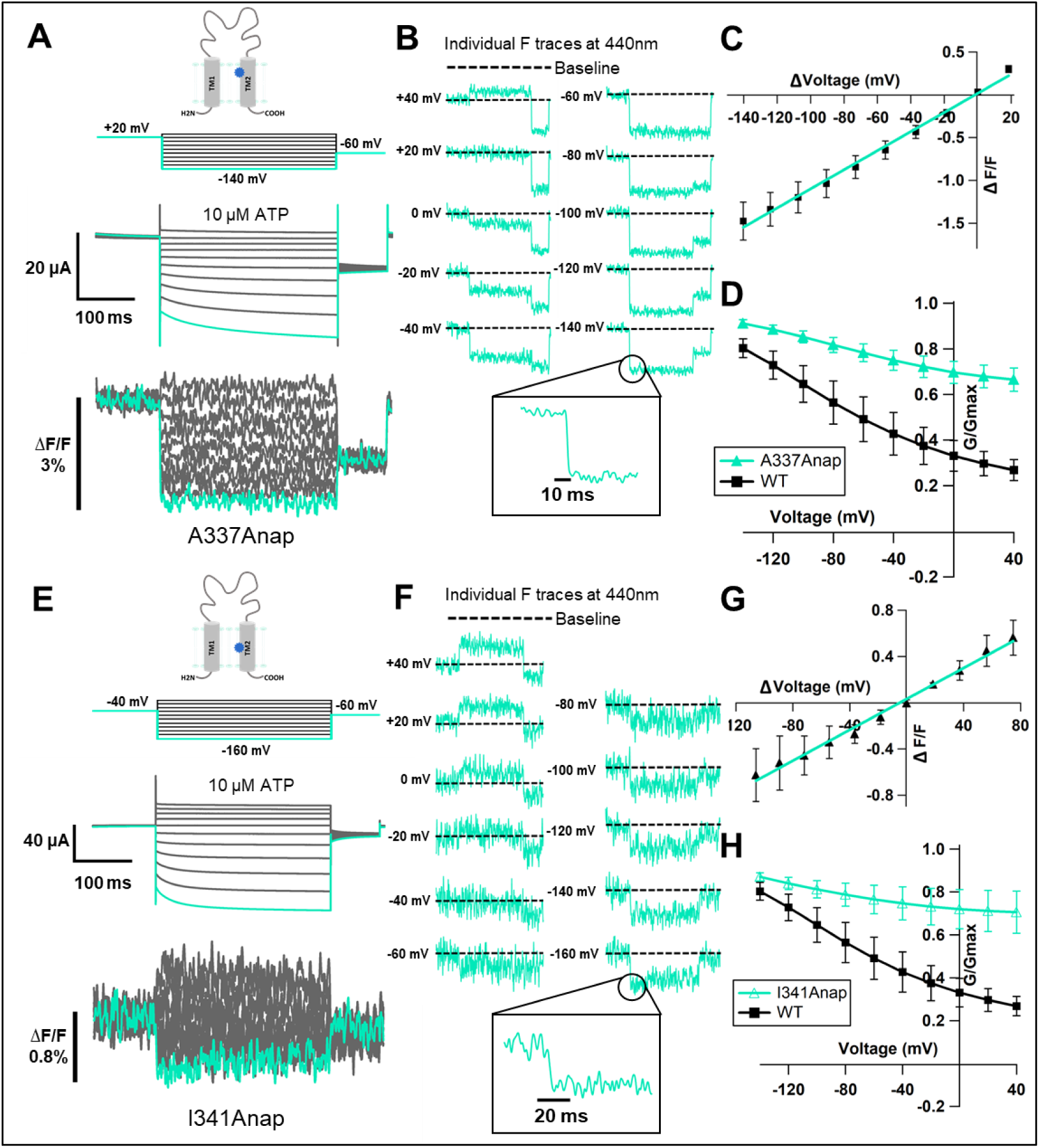
Voltage-clamp fluorometry of Anap-incorporated P2X2 receptor in the presence of 300 nM SIK inhibitor upon ATP and voltage stimuli. The focused electric field converged at A337 and I341 in TM2, throughout P2X2 ATP- and voltage-dependent gating. (**A**) Representative current traces and fluorescence signal of VCF recordings at A337, with 300 nM SIK inhibitor treatment, in the presence of 10 μM ATP (ΔF/F= 1.5%±0.2 at 440 nm, n=8). (**B**) Individual fluorescence traces during each voltage step at 440 nm. Inset shows fluorescence changes exhibiting fast kinetics in ms range. (**C**) F-V relationship showed a linear voltage-dependence. Each X-axis for F-V relationship is ΔV from the holding potential. (**D**) G-V relationship comparison between A337Anap (turquoise filled triangle) and wildtype (black filled square) for 10 μM ATP (n=8). Normalization was done based on the maximum conductance in the same concentration of ATP (10 μM) for each construct. (**E**) Representative current traces and fluorescence signal of VCF recordings at I341, with 300 nM SIK inhibitor treatment, in the presence of 10 μM ATP (ΔF/F= 0.6%±0.2 at 440 nm, n=3). (**F**) Individual fluorescence traces in each voltage step at 440 nm. Inset shows fluorescence changes also exhibiting fast kinetics in ms range. (**G**) F-V relationship showed a linear voltage-dependence. Each X-axis for F-V relationship is ΔV from the holding potential. (**H**) G-V relationship comparison between I341Anap (turquoise open triangle) and wildtype (black filled square) for 10 μM ATP (n=3). Normalization was done based on the maximum conductance in the same concentration of ATP (10 μM) for each construct. All error bars are ± s.e.m centered on the mean. (**Source Data 1 Figure 3**)

Electrochromic effect is known as a shift in the fluorophore emission spectrum due to the interaction between two components: the fluorophore electronic state and the local electric field (Bublitz & Boxer, 1997; Klymchenko & Demchenko, 2002; Dekel et al., 2012). It has two distinctive characteristics: (1) fast kinetics of fluorescent change (ΔF_Fast_); (2) linear voltage-dependence of the F-V relationship (Asamoah et al., 2003; Klymchenko et al., 2006). The electrochromic effect in some voltage-sensitive dyes is used to directly detect the change of membrane potential by attaching the dye to the cell membrane. If the fluorophore is directly attached in a site-specific manner within ion channels / receptors as shown by studies in the *Shaker* B K^+^ channel (Asamoah et al., 2003) and M_2_ muscarinic receptor (Dekel et al., 2012), the detection of electrochromic effect implies that there is a convergence of the electric field at the position where the fluorophore is attached. Thus, the observed fluorescence change at the position of A337 in the P2X2 receptor was explained to be due to the electrochromic effect, indicating that there is a focused electric field at A337 in the TM2 domain.

We noted that the G-V relationship for this mutant showed that a large fraction of the channel is already open, even at depolarized potentials, in 10 μM ATP, compared to wildtype (**Fig. 3D**), because of the high density of the expressed channel shown by a rather large current amplitude (> 20 μA). A previous study showed that P2X2 channel properties are correlated with expression density (Fujiwara & Kubo, 2004). In the case of lower expression levels, A337Anap showed a phenotype like wildtype. For the purpose of VCF experiments, however, a high expression level is needed to observe a detectable fluorescence change, and thus we needed to use oocytes with high expression, resulting in a lesser fraction of voltage-dependent activation. Nonetheless, we could still observe a weak voltage-dependent relaxation during hyperpolarization, and thus this fluorescence change still reflects an event occurring at or around the position of A337 when the receptor senses the change in membrane voltage.

Similarly, the application of 300 nM SIK inhibitor resulted in a clearer and more frequent Anap ΔF/F at the position of I341 in the TM2 (ΔF/F= 0.6%±0.2 at 440 nm, n=3, **Fig. 3E**) upon voltage step application in 10 μM ATP. The fluorescence intensity changes also occurred almost instantaneously in less than 5 ms (**Fig. 3F**). The ΔF/F – V relationship of I341Anap upon voltage step pulses in the presence of 10μM of ATP, from +40 mV to −160 mV with a holding potential at −40 mV, also showed a linear voltage-dependence (y = 0.007x + 0.03; R² = 0.99, n=3, **Fig. 3G**). Thus, ΔF observed at the position of I341 in the TM2 domain also did not correlate with hyperpolarization-induced conformational change. The changes were thought to be due to a phenomenon similar to that observed at the position of A337, which is related to electrochromic effect. The G-V relationship of this mutant in the presence of 10 μM ATP was not different from that of A337Anap, as shown in **Fig. 3H**. Taking these results together, the observed fluorescence intensity changes at I341 and A337 in the TM2 domain is best explained by an electric field convergence close to both positions which could be critical for the complex gating of the P2X2 receptor.

### Fluorescence change of Anap at A337 upon voltage change was observed also in 0 ATP condition and was [ATP]-dependent

To ensure that the fluorescence changes observed at A337 upon voltage change were not due to a change of ion flux, as in the case of the K2P K^+^ channel (Schewe et al., 2016), recording with the application of voltage step pulses was performed in the absence of ATP. In the same cells, VCF recordings were performed by applying voltage step pulses in the absence of ATP and then in the presence of 10 μM ATP.

When the voltage step pulses were applied in the absence of ATP, fluorescence changes were observed (ΔF/F= 1.9%±0.4 at 440 nm, n=4, **Fig. 4A – C**). The changes also exhibited fast kinetics and a linear voltage-dependence. ΔF/F in the absence of ATP was larger than that in the presence of 10 μM ATP (ΔF/F= 0.7%±0.1 at 440 nm, n=4, **Fig. 4A – C**). Thus, the focused electric field at the position of A337 is stronger in the absence of ATP than in the presence of 10 μM ATP.

**Figure 4.**
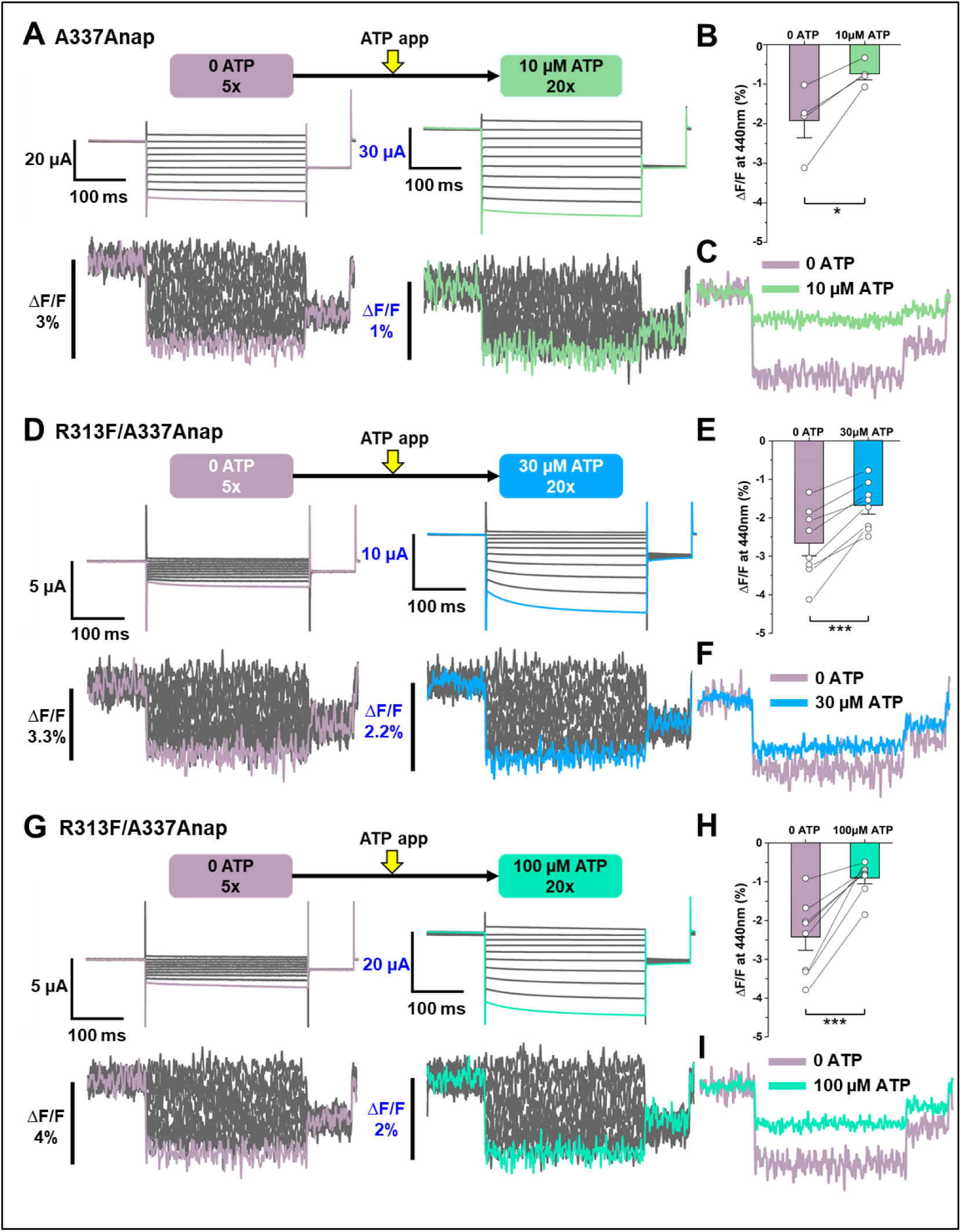
Voltage-clamp fluorometry of Anap-labeled P2X2 at A337 in TM2 evoked by hyperpolarization in the absence and presence of ATP. Anap fluorescence changes at A337 were also observed even in the absence of ATP upon hyperpolarization. (**A**) Representative current traces and fluorescence signal of VCF recordings at A337 in the absence of ATP (ΔF/F= 1.9%±0.4 at 440 nm, n=4) and in the presence of 10 μM ATP (ΔF/F= 0.7%±0.1 at 440 nm, n=4), from the same cell. (**B**) Comparison of the fluorescence changes in the absence and in the presence of 10 μM ATP (* p≤0.05, p=0.029, paired t-test, n=4) (**C**) Superimposed fluorescence traces at −140 mV, in 0 ATP (light purple) and 10 μM ATP (light green), from the same cell. (**D-I**) An additional R313F mutation was introduced to lower the basal activity of A337Anap and stabilize the closed state. (**D**) Representative current traces and fluorescence signal of VCF recordings of A337/R313F in the absence of ATP (ΔF/F= 2.6%±0.3 at 440 nm, n=8) and in the presence of 30 μM ATP (ΔF/F= 1.7%±0.2 at 440 nm, n=8) from the same cell. (**E**) Comparison of the fluorescence changes in the absence and in the presence of 30 μM ATP (***, p≤0.001, p=0.00045, paired t-test, n=8) (**F**) Superimposed fluorescence traces at −140 mV, in 0 ATP (light purple) and 30 μM ATP (blue), from the same cell. (**G**) Representative current traces and fluorescence signal of VCF recordings of A337/R313F in the absence of ATP (ΔF/F= 2.4%±0.3 at 440 nm, n=8) and in the presence of 100 μM ATP (ΔF/F= 0.9%±0.1 at 440 nm, n=8). (H) Comparison of the fluorescence changes in the absence and in the presence of 100 μM ATP (*** p≤0.001, p=0.0005, paired t-test, n=8). (I) Superimposed fluorescence traces at −140 mV in 0 ATP (light purple) and 100 μM ATP (turquoise), from the same cell. All error bars are ± s.e.m centered on the mean. (**Source Data 1 Figure 4**)

However, the A337Anap mutant showed a high basal activity, even in the absence of ATP, when the expression level was high. As observed in the current traces in no ATP, some of the channels expressed were already open (**Fig. 4A**). Thus, the fluorescence changes in 0 ATP observed in the above experiments might just represent the focused electric field in the open state. To record ΔF in the closed state with little current in no ATP, an additional mutation was introduced which suppresses the basal activity by stabilizing the closed state.

The extracellular linker plays important roles in transmitting the signal from the ATP binding pocket (ECD domain) to the TM domains (Keceli & Kubo, 2014). A mutation of R313 at the extracellular linker in β-14, which directly links the ATP binding site in the ECD domain with the TM2 domain to phenylalanine or tryptophan stabilized the closed state of the P2X2 receptor, as seen in the G-V relationship in 100 μM ATP (**Fig. 4 Supplementary Figure 2 A – D**). This mutation was introduced on top of A337Anap (A337Anap/R313F or A337Anap/R313W) to determine whether the focused electric field is present at the position of A337 even when the channel is mostly closed in 0 ATP.

Both results from VCF recording of A337Anap/R313F (**Fig. 4D – I**) and A337Anap/R313W (**Fig. 4—figure supplement 1 F – K**) confirmed that the focused electric field is present at A337 even when the channel is mostly closed. VCF recording in the absence of ATP for A337Anap/R313F showed a remarkable ΔF/F with mostly closed channels when voltage step pulses were applied (ΔF/F= 2.6%±0.3 at 440 nm, n=8; **Fig. 4D – F**). 30 μM ATP application resulted in smaller ΔF/F than in 0 ATP (ΔF/F= 1.7%±0.2 at 440 nm, n=8; **Fig. 4D – F**). These results confirmed that the focused electric field at the position of A337 is stronger in the absence of ATP than in the presence of ATP.

It is of interest to know whether or not the concentration of ATP affects the focused electric field at A337. Therefore, a higher concentration of ATP (100 μM) was tested for the same series of experiments. Fluorescence changes were again larger in the absence of ATP (ΔF/F= 2.4%±0.3 at 440 nm, n=8 **Fig. 4G – I**) and smaller in the presence of 100 μM ATP (ΔF/F= 0.9%±0.1 at 440 nm, n=8 **Fig. 4G – I**). ΔF/F was shown to become smaller with an increase in [ATP], by comparing ΔF/F in the presence of 30 μM and 100 μM ATP.

Similar series of experiments were also performed using A337Anap/R313W construct (**Fig. 4—figure supplement 1 F – K**), and similar phenotypes were observed. Taken together, these results show that the focused electric field at A337 is [ATP]-dependent and stronger in the absence of ATP, suggesting that the rotation of TM1 upon ATP binding (**Fig. 8**) would tighten the space surrounding A337 making the electric field more converged.

### Hyperpolarization-induced structural rearrangements were detected at or around A337 in TM2 upon the additional mutation of K308R

Upon ATP binding, the P2X receptor undergoes major structural rearrangements which result in transitions from closed to open state, with remarkable alterations in the three regions: ATP binding site, extracellular linker, which links ECD to TM domains, and TM domains (Kawate et al., 2009; Hattori & Gouaux, 2012; Mansoor et al., 2016). There is a possibility that the P2X2 receptor could undergo relatively minor but important structural rearrangements in response to hyperpolarization of the membrane voltage after the overall structure is altered greatly by the binding of ATP. A fraction of a slow fluorescence intensity change and non-linear ΔF/F – V could not be detected by the VCF experiments so far. This might be due to less clear voltage-dependent activation in high expression oocytes, with a significant activity even at depolarized potentials (e.g. **Fig. 3D, H**). Thus, an additional mutation which shows remarkable voltage-dependent activation, even in high expression conditions, is needed.

We then tested this possibility by introducing a K308R mutation on top of A337Anap. This charge-maintaining mutation, K308R, is shown to make the voltage-dependent activation more prominent, i.e. it is least active at depolarized potentials, even in high-expression oocytes, and it also accelerates the activation kinetics of P2X2 upon voltage stimuli (Keceli & Kubo, 2009). K308 is a conserved residue located in the ATP binding site. It was shown to be not only important for ATP binding (Ennion et al., 2000; Jiang et al., 2000; Roberts et al., 2006) but also for the conformational change associated with channel opening (Cao et al., 2007). If the voltage-dependent activation is more prominent even in high expression cells for VCF experiments, there is a possibility that we might be able to detect the fluorescence intensity change associated with the voltage-dependent gating.

VCF recording of K308R/A337Anap was performed in the presence of 300 μM ATP, while a voltage-step from +40 mV to −160 mV, with a holding potential of +20 mV, was applied. A high concentration of ATP was applied because K308R/A337Anap has a lower sensitivity to ATP. Hyperpolarization elicited fluorescence signals which consist of two components, a very fast decrease (ΔF_Fast_/F) and a slow increase (ΔF_Slow_/F), until it reached the steady-state (ΔF_Steady-state_/F) (**Fig. 5A, B**). Plots of the F-V relationship at the end of the recording time course (at the steady-state), showed that ΔF/F – V consists of mixed components, a linear component and a non-linear component (**Fig. 5C**). The presence of the two components suggests that they might result from two different mechanisms. The F-V relationship of ΔF_Fast_/F showed a linear voltage-dependence, which is similar to the F-V for A337Anap alone, which was generated from the electrochromic signal (**Fig. 3C, Fig. 5D**). In contrast, the F-V relationship of ΔF_Slow_/F showed a non-linear voltage-dependence. The F-V and G-V relationships of the slow component overlap very well (**Fig. 5E**), showing that the slow F change reflects the hyperpolarization-induced structural rearrangements that occur at or around the position of A337.

**Figure 5.**
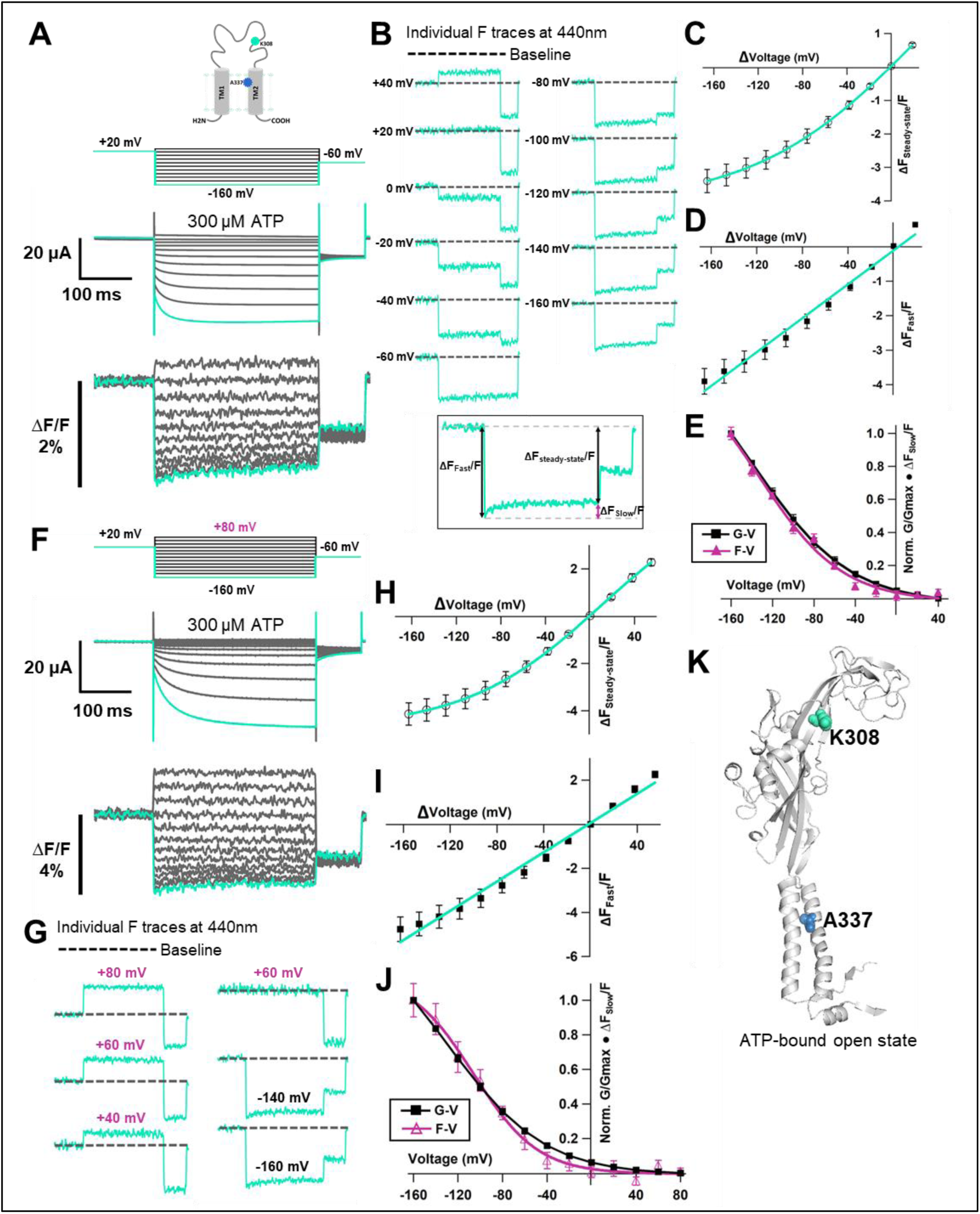
Voltage-clamp fluorometry of Anap-labeled P2X2 at A337 in TM2 with the additional mutation of K308R evoked by hyperpolarization in the presence of ATP. (**A**) Representative current traces and fluorescence signal of VCF recordings of K308R/A337Anap with 300 nM SIK inhibitor treatment in the presence of 300 μM ATP, from +40 mV to −160 mV with a holding potential of +20 mV (ΔF_Steady-state_/F= 3.4%±0.3 at 440 nm, n=8). (**B**) Individual fluorescence traces at each voltage step. Inset shows that the fluorescence signal of K308R/A337Anap consists of two components, instantaneous downward change (ΔF_Fast_/F) and slow upward change (ΔF_Slow_/F). (**C**) F-V relationship of the mixed component (ΔF_Steady-state_/F) was calculated from the last 50 ms of fluorescence signal. Component of ΔF_Steady-state_/F is shown in inset of (B). F_Steady-state_ – V relationship shows that it consists of only a linear component at depolarized potentials, and there are mixed components at hyperpolarized potentials. (**D**) F_Fast_ – V relationship was taken from the first 5 ms of the fluorescence signal. F_Fast_ – V relationship showed almost linear voltage-dependence (ΔF_Fast_/F=3.9%±0.4 at 440nm, n=8). (**E**) Comparison of F_Slow_ – V and G-V relationships. Purple filled triangle trace shows F_Slow_ – V relationship extracted from the fluorescence traces depicted in inset (B), as shown by purple arrow, from the equation ΔF_steady-state_/F = ΔF_fast_/F + ΔF_slow_/F. Normalization was done based on the maximum ΔF_slow_/F (at −160 mV). Black filled square trace shows G-V relationship in the presence of 300 μM ATP. Normalization was done based on the maximum conductance in the same concentration of ATP (300 μM). (**F**) Representative current traces and fluorescence signal of VCF recordings of K308R/A337Anap with 300 nM SIK inhibitor treatment, in the presence of 300 μM ATP, at more depolarized potentials from up to +80 mV to −160 mV, with a holding potential of +20 mV (ΔF_steady-state_/F=4.1%±0.5 at 440 nm, n=5). (**G**) Individual fluorescence traces at each depolarized voltage step and some hyperpolarized voltage steps. (**H**) F_Steady-state_ – V relationship further confirms that it consists of a linear component and a slow component only generated upon hyperpolarization. (**I**) F_Fast_ – V relationship shows almost linear voltage-dependence (ΔF_Fast_/F=4.7%±0.5 at 440nm, n=5). (**J**) Comparison of F_Slow_ – V and G-V relationships. Purple open triangle trace shows F_Slow_ – V relationship extracted from the fluorescence traces depicted in (F). Normalization was done based on the maximum ΔF_slow_/F (at −160 mV). Black filled square trace shows G-V relationship in the presence of 300 μM ATP. Normalization was done based on the maximum conductance in the same concentration of ATP (300 μM). All error bars are ± s.e.m centered on the mean. (**K**) Side view structure of the position of K308 and A337 in the ATP-bound open state. (**Source Data 1 Figure 5**)

Next, we examined whether ΔF_Slow_/F is indeed generated only at hyperpolarized potential to confirm that this is evidence of voltage-dependent structural rearrangements during P2X2 receptor complex gating. We performed VCF recordings by applying step pulses from up to +80 mV to −160 mV, with a holding potential of +20 mV. The F-V relationship in the steady-state showed a mixed signal. This set of recordings showed that at more depolarized potentials the fluorescence signal consists only of a linear component (**Fig. 5F – H**). Separation of the mixed fluorescence signal also resulted in a rapidly changing linear F-V for ΔF_Fast_/F (**Fig. 5I**) and a non-linear F-V for ΔF_Slow_/F (**Fig. 5J**) with no slow component from +80 mV to 0 mV.

The results further confirm that the slow rise in K308R/A337Anap fluorescence signal reflects the structural rearrangements at or around the position of A337 in response to the change in membrane voltage.

### Fluorescence signal changes at A337Anap/K308R exhibited only the fast component in the absence of ATP and showed two components in the presence of ATP

We also examined whether the non-linear component of the K308R/A337Anap fluorescence signal was abolished in the absence of ATP. We then performed VCF recordings of the same cell by applying voltage steps in the absence of ATP and in the presence of 300 μM ATP. In the absence of ATP, the fluorescence signal consisted of only one component, the fast component (ΔF_Fast_/F, **Fig. 6A**). The F-V relationship for this fast component was linear and is thought to be derived from the electrochromic phenomenon, showing that A337 is located in the focused electric field (**Fig. 6C**).

**Figure 6.**
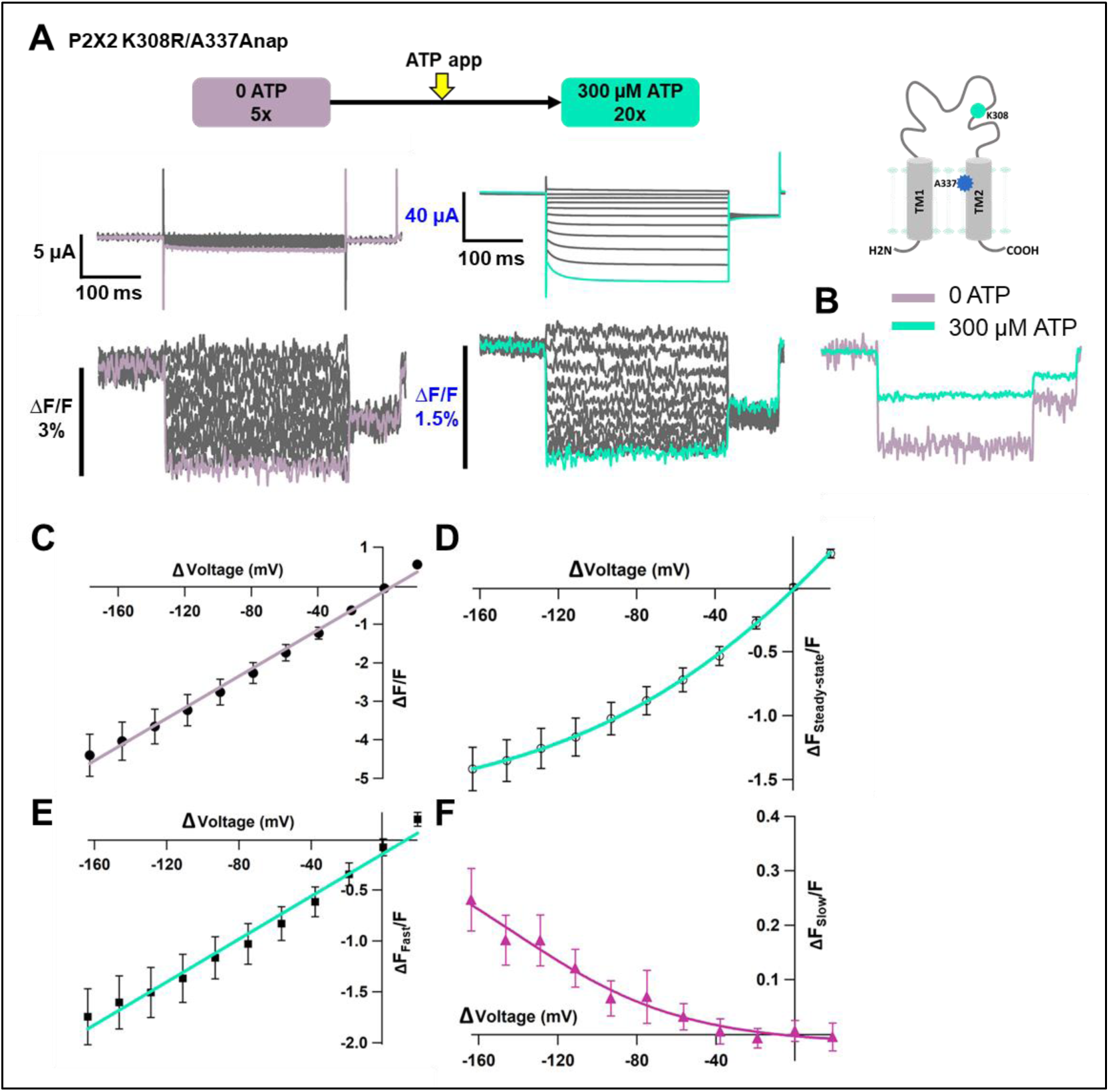
Voltage-clamp fluorometry of Anap-labeled P2X2 at A337 in TM2 with the additional mutation of K308R evoked by hyperpolarization in the absence and presence of ATP. Fluorescence signal changes at K308R/A337Anap exhibited only a fast component in the absence of ATP and consisted of two components in the presence of ATP. (**A**) Representative current traces and fluorescence signal of VCF recordings of K308R/A337Anap in the absence of ATP (ΔF/F= 4.4%±0.5 at 440 nm, n=6) and in the presence of 300 μM ATP (ΔF_Steady-state_/F=1.4%±0.2 at 440 nm, n=6), from the same cell. (**B**) Superimposed fluorescence traces at −160 mV in 0 ATP (light purple) and 300 μM (turquoise). (**C**) F-V relationship, in the absence of ATP, taken from the last 100 ms of the fluorescence signals shows a linear voltage-dependence (R² = 0.99); therefore, it has only the fast component (ΔF_Fast_/F). (**D**) F-V relationship, in the presence of 300 μM ATP, taken from the last 50 ms (ΔF_Steady-state_/F) of the fluorescence signals shows mixed components. (**E-F**) F-V relationship from two separate components of the fluorescence signal change, in the presence of 300 μM ATP. (**E**) F_Fast_ – V relationship (ΔF_Fast_/F=1.7±0.3 at 440nm, n=6) shows almost linear voltage-dependence (R² = 0.98). (**F**) F_Slow_ – V relationship (ΔF_Slow_/F=0.25±0.05 at 440nm, n=6). Each X-axis for the F-V relationship is ΔV from the holding potential. All error bars are ± s.e.m centered on the mean. (**Source Data 1 Figure 6**)

Subsequently, when the voltage step pulses were applied in the presence of 300 μM ATP, the slow component could be observed (**Fig. 6A, D**). The F-V relationship in the steady-state showed a mixture of the two components (**Fig. 6D**). Separation of this mixed component resulted in a linear F-V for the fast component (**Fig. 6E**) and a non-linear F-V for the slow component (**Fig. 6F**), which is consistent with the previous experiments. Additionally, consistent results were also obtained in terms of the fluorescence intensity change of the fast component. ΔF_Fast_/F in the absence of ATP was larger than in the presence of ATP (ΔF_Fast_/F=4.4%±0.5 at 440 nm, n=6 and ΔF_Fast_/F=1.7%±0.3 at 440 nm, n=6; **Fig. 6B**). Taken together, these results further show that the slow component of the fluorescence intensity changes reflects the structural rearrangements of the P2X2 receptor upon complex gating which depends on both [ATP] and voltage.

### A337 in TM2 might interact with F44 in TM1 to stabilize the open state of the P2X2 receptor

The electric field convergence at A337 and I341 and the voltage-dependent conformational changes at or around A337 could provide us with a clue to understand the mechanism of the complex gating of the P2X2 receptor. The existence of a strong electric field supports the possible location of a key residue which is responsible for the voltage sensing (Asamoah et al., 2003; Dekel et al., 2012). Thus, various single amino acid mutations were introduced at the position of A337, and their electrophysiological properties were analyzed, focusing on the [ATP]-dependent and voltage-dependent gating properties, to see whether or not this amino acid plays an important role in the P2X2 complex gating (**Fig. 7A – B**).

**Figure 7.**
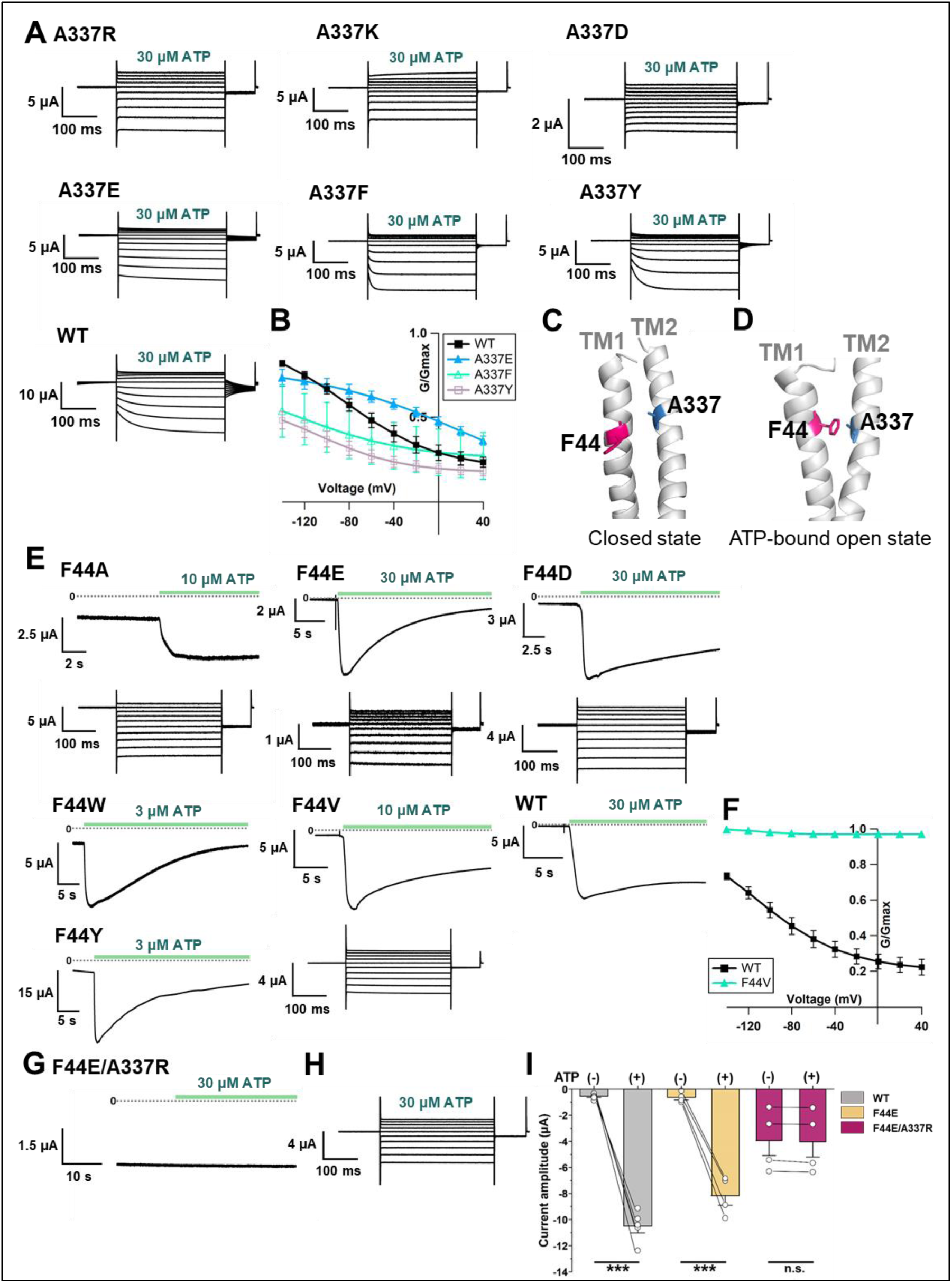
Effects of mutations at A337 in TM2 and F44 in TM1 on P2X2 receptor ATP- and voltage-dependent gating. (**A**) Representative current traces of single amino acid mutants at the position of A337 in the presence of 30 μM ATP in response to voltage step pulses from +40 mV to −140 mV, with a holding potential of −40 mV (A337R, A337K, A337D, A337E, A337F, A337Y, and WT; respectively). (**B**) Comparison of G-V relationships between WT (black filled square), A337E (blue filled triangle), A337F (turquoise open triangle), and A337Y (purple open square) for 30 μM ATP (n=3), from tail current analysis at −60 mV. Normalization was done based on the maximum conductance in the highest [ATP] (300 μM) for each construct. (**C, D**) Side view structure of the position of F44 (magenta) and A337 (blue) in the closed (**C**) and ATP-bound open (**D**) state, respectively. (**E**) Representative current traces of single amino acid mutants at the position of F44 upon application of various [ATP] (F44A, F44W, F44Y, F44E, F44D and WT; respectively; n=3-6 for each mutant). (**F**) G-V relationship comparison between WT (black filled square) and F44V (turquoise filled triangle) for 10 μM ATP (n=3), showing that this mutant was equally active at all recorded voltages and was far less sensitive to voltage than wildtype. Normalization was done based on the maximum conductance in the highest [ATP] (300 μM) for each construct. (**G, H**) Representative current traces of F44E/A337R upon ATP (**G**) and voltage (**H**) application. (**I**) Comparison of current amplitude of WT, F44E, and F44E/A337R before and after ATP application (*** p≤0.001, p=0.00007 for WT and p= 0.00095 for F44E, paired t-test, n=4-5). All error bars are ± s.e.m centered on the mean. (**Source Data 1 Figure 7**)

Mutations to A337R, A337K, and A337D had severe effects. When the voltage step pulses were applied in 30 μM ATP, these mutants almost lacked voltage sensitivity. A337E, A337Y, and A337F showed a voltage sensitivity with various activation kinetics. The most striking changes were observed in A337Y and A337F. The activation evoked by a voltage step was clearly different from wildtype, whereas the A337E mutation had a less severe effect (**Fig. 7A**). G-V relationships in 30 μM ATP for mutants and wildtype were analyzed (**Fig. 7B**). Normalization was done based on the maximum conductance at the highest ATP concentration (300 μM) from each construct. Here we could also see that the mutants of A337Y and A337F preferred to stay in the closed state. As the activation kinetics and the voltage dependence were altered by the introduction of mutation at A337, this position was shown to be critical for the P2X2 receptor complex gating.

Next, we aimed to identify the counter-part in the TM1 domain with which A337 might have an interaction during the complex gating. Based on the homology modelling of *r*P2X2 in the closed and ATP-bound open states from *h*P2X3 crystal structure data (PDB ID: 5SVJ, 5SVK, respectively) (Mansoor et al., 2016), F44 in the TM1 domain was shown to rotate and move towards A337 upon ATP binding (**Fig. 7C, D**). Various single amino acid mutations were then introduced at F44 and their [ATP]-dependent and voltage-dependent gating was analyzed (**Fig. 7E, F**).

The F44A mutation strikingly changed the gating. It showed a relatively high basal current in the absence of ATP and further responded to ATP application. Voltage-dependent gating was also changed, as seen in the lack of tail current, showing that this mutant might have a constitutive activity with rectified permeation properties. Mutation to positively charged residues (F44R, F44K) resulted in a non-functional channel and/or a very low expression level, as the recording on day 4 did not evoke any response to the highest concentration of ATP used in this study (300 μM). Mutation to negatively charged residues (F44E, F44D) and aromatic residues (F44Y, F44W) remarkably changed the ATP-evoked response (**Fig. 7E**).

All four mutants still opened upon the application of ATP but current decay in the continuous presence of ATP appeared to be faster than wildtype.

F44 is conserved only in P2X2 and P2X3. Other subtypes of P2X receptor, like P2X1, P2X4, P2X6, and P2X7, except P2X5, have valine at the corresponding position (Kawate et al., 2009). Thus, the F44V mutation was also introduced. 10 μM of ATP could activate F44V but resulted in faster current decay than wildtype. Voltage step pulses were applied during the course of current decay because there was no clear steady-state (**Fig. 7E**). Nonetheless, it could still be observed how the mutation at F44V changed the voltage-dependent gating. The G-V relationship of F44V in 10 μM ATP showed that this mutant was far less sensitive to voltage than wildtype (**Fig. 7F**). Taken together, the results of the mutations introduced at position F44 showed that this residue is critical for the proper ATP- and voltage-dependent gating of the P2X2 receptor.

Additionally, as the single amino acid mutations at both A337 and F44 altered the gating of P2X2, it is of interest to know whether the introduction of swapped mutations into A337/F44 would rescue the wildtype phenotype. The phenotype of F44A/A337F was similar to F44A and the wildtype phenotype was not rescued (**Fig. 7—figure supplement 1**).

It is possible that an interaction between A337 and F44 could not be properly formed in the swapped mutant.

Next, an artificial electrostatic bridge was introduced between A337 and F44 to prove that the interaction between the two residues is critical in the ATP-bound open state. Various paired electrostatically charged residues were introduced into A337 and F44, in order to see if the artificial electrostatic bridge could be formed. The F44E/A337R pair showed a constitutive activity. This double mutant was already open before ATP application and didn’t show any response to ATP application (**Fig. 7G**). When voltage step pulses were applied, this mutant lacked sensitivity to voltage with a rectified permeation property, as seen by the total lack of tail currents (**Fig. 7H**). Additionally, the comparison of the current amplitude before and after ATP application showed that F44E/A337R is already open before ATP application (**Fig. 7I**). The results showed that A337 in the TM2 domain might interact with F44 in TM1 to stabilize the open state of the P2X2 receptor.

Based on the results from VCF recording, mutagenesis experiments, and the homology modeling of *r*P2X2 in the open state upon ATP binding, it was shown that F44 moves into close proximity to the converged electric field at A337 and I341 (**Fig. 8B, C**). In the presence of ATP, voltage-dependent conformational changes occur possibly at or around the position of A337 and F44, giving influence to the interaction between A337 and F44, which is critical for stabilizing the open state. Results of this study show that the origin of the voltage-dependent gating of P2X2 in the presence of ATP is possibly the voltage dependence of the interaction between A337 and F44 in the converged electric field.

**Figure 8.**
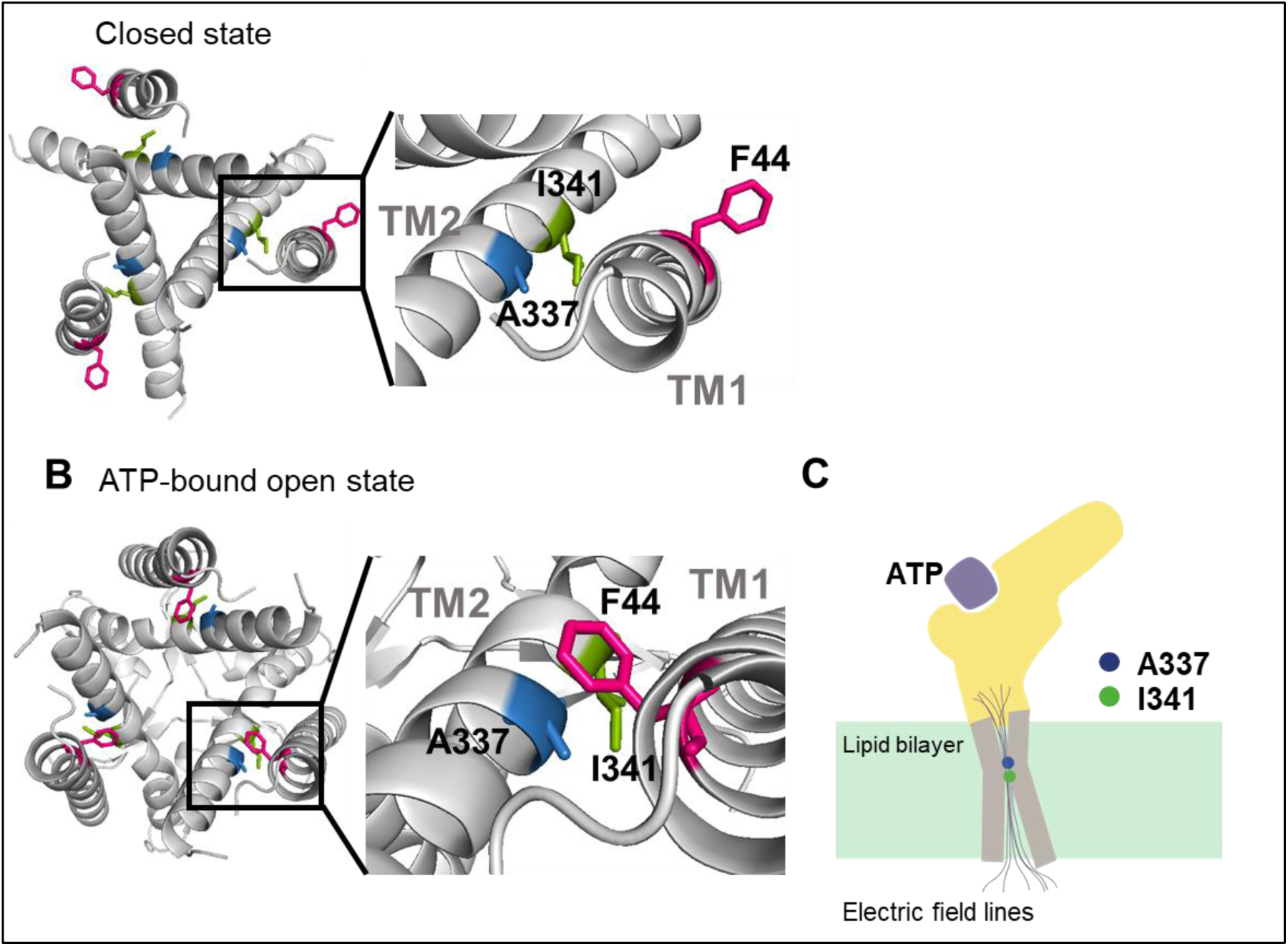
Proposed initiation mechanisms of P2X2 receptor complex gating. **(A, B)** Top view structure of P2X2 receptor in the closed (A) and ATP-bound open state (B). Depicted are the proposed initiation mechanisms of P2X2 receptor complex gating as follows. (1) The electric convergence at A337 and I341 (2) F44 moves towards A337 in TM2 domain upon ATP binding (3) Hyperpolarization-induced structural rearrangements around A337 in TM2 (4) The interaction between A337 and F44 in the ATP-bound open state is thought to be under the influence of the converged electric field. **(C)** A schematic illustration of the focused electric field at A337 and I341. Ion permeation pathway is not depicted in this scheme.

## Discussion

The present study aims at defining the roles of the TM domains of the P2X2 receptor in the complex gating by [ATP] and voltage, using VCF with a genetically incorporated fUAA probe, named Anap, and a mutagenesis study. The following findings were obtained.

### Detection of fast F changes with a linear voltage-dependence at A337 and I341

We analyzed 96 mutants by VCF and detected voltage-dependent ΔF_Fast_/F change at the position of A337 and I341 in TM2. It was very fast and showed a linear voltage-dependence in the recorded voltage range. The change could be well interpreted to be due to an electrochromic effect, indicating that there is an electric field convergence at both positions, which are located adjacent to each other.

An electrochromic signal is an intrinsic property exhibited by voltage-sensitive fluorescent dyes or electrochromic probes to directly detect transmembrane potentials (Loew, 1982; Zhang et al., 1998). By standard use of electrochromic probes in a lipid bilayer, it is hard to sense the electrical potential that directly acts on the voltage-sensing machinery of membrane proteins (Asamoah et al., 2003). This is because the local electric field at a certain position in the lipid bilayer is not steep enough. On the other hand, previous VCF studies on the *Shaker* K^+^ channel, using modified electrochromic probes (Asamoah et al., 2003), and on the M_2_ muscarinic receptor, using TMRM (Dekel et al., 2012), showed that an electrochromic signal could also be observed when the fluorophore is directly attached to a specific position within the ion channel / receptor. These studies stated that this phenomenon did not report conformational changes of the protein at a specific position where the fluorophore was attached, but rather implied that there is an electric field convergence if the electrochromic signal is observed only at positions adjacent to each other (Asamoah et al., 2003; Dekel et al., 2012). This observed electrochromic signal might support the possible location of a voltage sensor (Asamoah et al., 2003; Dekel et al., 2012). Further studies are certainly required to prove this possibility.

An almost linear F-V relationship which might originate from the electrochromic signal was also reported from VCF studies in a canonical VSD-containing membrane protein named hTMEM266 labeled with MTS-TAMRA. The observed ΔF_Fast_/F was, however, explained rather differently. Even though the ΔF_Fast_/F was observed at most of the introduced positions located in the S3-S4 linker and the top of the S4 segment, it was stated that ΔF_Fast_/F was not due to a direct electrochromic effect but instead was associated with rapid voltage-dependent conformational changes on a μs time scale (Papp et al., 2019). In the case of hTMEM266, it is hard to surmise that the fast change detected at many positions is due to electrochromic effect, because it suggests an unlikely possibility that the electric field is converged at various positions. Conversely, in the P2X2 receptor, there were only two adjacent positions which exclusively showed ΔF_Fast_/F and a linear F-V relationship.

In the hTMEM266 study, it was also a concern whether TAMRA-MTS could report an electrochromic signal, because there was not any previous finding to explain this case. There was also no report of electrochromic signals recorded using Anap as fluorophore to date. Anap has only been reported as an environmentally sensitive fluorophore (Lee et al., 2009; Chatterjee et al., 2013). None reported that Anap is an electrochromic fluorophore, unlike the case of the modified fluorophore used in *Shaker* Kv studies, which has been reported to have electrochromic properties (Zhang et al., 1998; Asamoah et al., 2003). On the other hand, studies on the M_2_ muscarinic receptor did not discuss TMRM fluorophore properties, but still concluded that the observed fast F change with linear F-V originated from the electrochromic signal (Dekel et al., 2012). Even though other possibilities could still remain, the most straightforward explanation to interpret the results observed in this study is that the very fast and linearly voltage-dependent fluorescence changes of Anap at A337 and I341 are associated not with the conformational changes of the P2X2 protein but presumably with the electrochromic signal. Consequently, the results show that there is an electric field convergence at these positions which could give us a clue about the possible location of the voltage sensor in the P2X2 receptor.

We observed that ΔF_Fast_/F changed with voltage in both the closed and ATP bound-open states, implying the presence of the focused electric field in both states at the position of A337. The focused electric field was more prominent in the absence of ATP. Some Cys accessibility studies were performed on the P2X2 receptor in the TM2 domain, to analyze the ATP-evoked gating mechanism (Li et al., 2008; Kracun et al., 2010; Li et al., 2010). A337 Cys mutants were first reported to be not modified by MTSET both in the presence or absence of ATP, indicating that these residues are not involved either in the pore lining region in the open state or in the gate of P2X2 (Li et al., 2008). Meanwhile in another study using Ag^+^, a smaller thiol-reactive ion with higher accessibility, A337C was modified both in the absence and presence of ATP (Li et al., 2010). These results suggest that a narrow water-phase penetrates down to this position, which is consistent with the results in this study that there is a focused electric field at A337.

### Detection of slow F change with non-linear voltage-dependence at A337 of K308R mutant

We obtained data supporting voltage-dependent conformational rearrangements occurring at or around the position of A337, by analyzing the mixed Anap fluorescence signal changes which contain both ΔF_Fast_/F and ΔF_Slow_/F in the presence of an additional mutation of K308R on top of A337Anap. K308 is located in the ATP binding site and was reported to be important not only for ATP binding but also for the gating of the P2X2 receptor (Ennion et al., 2000; Jiang et al., 2000; Roberts et al., 2006; Cao et al., 2007). In VCF analysis, a high expression level is needed to detect F changes successfully, overcoming the influence of the background fluorescence. However, high expression makes the P2X2 channel activate even in the absence of ATP and also even at depolarized potentials, i.e. the G-V is shifted to the depolarized potential, which makes the voltage-dependent activation upon hyperpolarization unclear. To overcome this problem, we introduced the K308R mutation, which shifts the G-V relationship in the hyperpolarized direction, with much reduced activity at depolarized potentials (Keceli & Kubo, 2009). By introducing the K308R mutation, we could observe voltage-dependent gating better and succeeded in recording the slow and voltage-dependent F change at A337 (**Fig. 5**).

In addition, the ΔF_Slow_ component was observed only at hyperpolarized potentials and in the presence of ATP (**Fig. 5F – J; Fig. 6**). Also, the F_Slow_ – V and G-V overlapped well, showing that ΔF_Slow_ reflects the hyperpolarization-induced structural rearrangements at or around the position of A337 (Fig. 5E; Fig. 5J). A337 in TM2 is indeed in the converged electric field, as shown by the linear F – V relationship of the ΔF_Fast_ component (**Fig. 5D**), supporting the notion that the main focus for the voltage-sensing mechanism in the P2X2 receptor lies at or around A337.

### Interaction between A337 in TM2 and F44 in TM1 in the converged electric field

The specific function of each transmembrane domain of the P2X receptor had been defined before the crystal structure was solved but the information as to the role of each TM in P2X2 voltage-dependent gating is limited. TM1 is shown to play a role in the binding-gating process, as mutations in this region alter the agonist selectivity and sensitivity of channel gating (Haines et al., 2001; Li et al., 2004; Stelmashenko et al., 2014). In contrast, TM2 plays an essential role in permeation (Nakazawa et al., 1998; Khakh & Egan, 2005) and gating (Li et al., 2008).

Mutations of A337 in the present study suggested that this position is critical for the complex gating, as mutation to A337F and A337Y altered the channel gating as well as the activation kinetics upon the application of ATP and voltage (**Fig. 7A – B**). The possible counter-part for A337 is most likely the F44 residue in TM1. Based on the homology modelling of P2X2, in the ATP-bound open state, F44 rotates and moves towards TM2, specifically into the proximity of A337 (**Fig. 7C – D**). Mutagenesis at the position of F44 showed the importance of F44 to maintain the open state in the presence of ATP (**Fig. 7E – F**). The artificial electrostatic bridge formation experiment of the F44E/A337R mutant (**Fig. 7G – I**) induced constitutive activity in the absence of ATP and at all recorded voltages, confirming the importance of the interaction for the maintenance of the activated state, and also showing the dynamic and presumably voltage-dependent interaction between A337 and F44 in the presence of ATP. The structural rearrangement at F44 is of very high interest, but F44Anap was not functional, further showing the critical role of F44.

There are several types of voltage-sensing mechanism in membrane proteins (Bezanilla, 2008): (1) charged residues, as in the case of canonical voltage-gated ion channels (Y. Jiang et al., 2003; Swartz, 2008) (2) side-chains that have an intrinsic dipole moment, such as Tyr, as in the case of the M_2_ muscarinic receptor (Ben-Chaim et al., 2006; Navarro-Polanco et al., 2011; Dekel et al., 2012; Barchad-Avitzur et al., 2016); (3) the α-helix, with its intrinsic dipole moment, and (4) cavities within the protein structure, filled with free ions. Based on our main findings, the interaction between A337 and F44 in the ATP-bound open state might be under the influence of the converged electric field (Fig. 8A – C). The findings also clearly demonstrate that there are voltage-dependent structural rearrangements in the proximity of A337 in TM2. At this point, the details of how the interaction contributes to the voltage sensing of P2X2 cannot be answered yet. Further structural dynamics analysis at the position of F44 will help to elucidate the detailed mechanism of the complex gating of the P2X2 receptor.

## MATERIALS AND METHOD

### Ethical approval

All animal experiments were approved by the Animal Care Committee of the National Institutes of Natural Sciences (NINS, Japan) and performed obeying its guidelines.

### Molecular biology

Wild type (WT) *Rattus norvegicus* P2X2 (*r*P2X2) receptor cDNA (Brake et al., 1994) was subcloned into the BamH1 site of pGEMHE. TAG or any single amino acid mutation and/or double mutations were introduced using a Quikchange site-directed mutagenesis kit (Agilent Technologies). The introduced mutations were confirmed by DNA sequencing. mMESSAGE T7 RNA transcription kit (Thermo Fisher Scientific) was used to transcribe WT and mutant *r*P2X2 cRNAs from plasmid cDNA linearized by Nhe1 restriction enzyme (Toyobo). The tRNA-synthetase/Anap-CUA encoding plasmid was obtained from addgene. Salt form of fUAA Anap was used (Futurechem).

*Ciona intestinalis* voltage-sensing phosphatase (*Ci*-VSP) with a mutation in the gating loop of the phosphatase domain (F401Anap) was used as a positive control (Sakata et al., 2016). mMESSAGE SP6 RNA transcription kit (Thermo Fisher Scientific) was used for cRNA transcription of *Ci*-VSP.

### Preparation of *Xenopus laevis* oocytes

0.15% tricaine (Sigma-Aldrich) was used as an anesthetic reagent for *Xenopus laevis* before surgical operation for isolation of oocytes. After the final collection, the frogs were humanely sacrificed by decapitation. Follicular membranes were removed from isolated oocytes by collagenase treatment (2 mg ml^−1^; type 1; Sigma-Aldrich) for 6.5 hours. Oocytes were then rinsed and stored in frog Ringer’s solution (88 mM NaCl, 1 mM KCl, 2.4 mM NaHCO_3_, 0.3 mM Ca(NO_3_)_2_, 0.41 mM CaCl_2_, 0.82 mM Mg_2_SO_4,_ and 15 mM HEPES pH 7.6 with NaOH) containing 0.1% penicillin-streptomycin at 17 °C.

### Channel expression and electrophysiological recording of *r*P2X2

*Xenopus* oocytes injected with 0.5 ng of WT *r*P2X2 cRNA and incubated for 2 days at 17 °C showed a high expression level phenotype of WT *r*P2X2 that has less voltage dependence than those of low expression level of P2X2 (I < 4.0 μA at −60 mV) (Fujiwara & Kubo, 2004). To achieve low expression level, oocytes were injected with 0.05 ng of WT *r*P2X2 cRNA and incubated for 1-2 days. For *r*P2X2 mutants, oocytes were injected with 0.5 ng – 2.5 ng of cRNA and incubated for 1-3 days, depending on the desired expression level.

Voltage clamp for macroscopic current recording was performed by using an amplifier (OC-725C; Warner Instruments), a digital-analogue analogue-digital converter (Digidata 1440, Molecular Devices), and pClamp10.3 software (Molecular Devices). In TEVC recording, borosilicate glass capillaries (World Precision Instruments) were used with a resistance of 0.2–0.5 MΩ when filled with 3 M KOAc and 10 mM KCl. P2X2 bath solution contained 95.6 mM NaCl, 1 mM MgCl_2_, 5 mM HEPES, and 2.4 mM NaOH at pH 7.35 – 7.45. Ca^2+^ was not included in the bath solution in order to avoid the inactivation of the receptor and secondary intracellular effects, e.g. activation of Ca^2+^ dependent chloride channel currents, (Ding & Sachs, 2000).

ATP disodium salt (Sigma-Aldrich) was prepared in various concentrations (1 μM, 3 μM, 10 μM, 30 μM, 100 μM, 300 μM, 1 mM, and 3 mM) by dissolving it in the bath solution. For recording using step-pulse protocols, ATP was applied in two ways, depending on the purpose of the experiments and the phenotype of the mutants. (1) Direct application using a motorized pipette (Gilson pipetman) which was set to exchange the whole bath solution with a ligand-based solution. 2000 μL (five times larger than the bath volume) of ligand-based solution was applied. (2) Perfusion of a recording chamber using a perfusion system set (ISMATEC pump). In both cases, overflowed bath solution was continuously removed using a suction pipette by negative air pressure. Oocytes were held at −40 mV and voltage step pulses were applied in the range from +40 mV to −140 mV. Tail currents were recorded at −60 mV to measure conductance-voltage (G-V) relationships. Recordings were performed at room temperature (24±2 °C).

### Expression of Anap incorporated *r*P2X2 and *Ci*-VSP

For functional expression of channels with incorporated Anap, 1.25 ng of cDNA encoding the tRNA synthetase/Anap-CUA pair was injected into the nucleus of defolliculated *Xenopus* oocytes located in the center of the animal pole (Kalstrup & Blunck, 2013). Oocytes were then incubated for 24 hours at 17 °C to allow tRNA transcription and synthetase expression. The subsequent step was performed with minimization of light exposure, which otherwise may have excited the fluorophore. Either 1.4 – 12.6 ng of *r*P2X2 cRNA or 8.2 ng of *Ci*-VSP cRNA in which the target site was mutated to a TAG codon, was co-injected with 23 nL of 1 mM Anap. Oocytes were incubated in frog Ringer’s solution (containing 0.1% penicillin-streptomycin) for 1-3 days (*r*P2X2) or 3-5 days (*Ci*-VSP) depending on the desired expression level. In the absence of either tRNA synthetase/Anap-CUA plasmid or fUAA Anap, no channel expression was detected in *r*P2X2 Anap mutants, confirming that functional channels are expressed only when they successfully incorporated fUAA.

### SIK inhibitor application

HG 9-91-01 / SIK inhibitor (MedChem Express) was dissolved in DMSO to make a stock solution of 10 mM and kept as aliquots at −80 °C. SIK inhibitor was diluted before use with RNase-free water (Otsuka) into certain concentrations for injection to oocytes. Various concentrations of SIK inhibitor were injected into oocyte nuclei to determine the most effective concentration to improve the optical recording of VCF-fUAA. SIK inhibitor was mixed and co-injected with either (1) tRNA synthetase/Anap-CUA plasmid (nuclear injection) or (2) cRNA + Anap (cytoplasmic injection). 300 nM was defined as the amount of the co-injected SIK inhibitor in the mixed solution. For instance, the actual concentration of SIK inhibitor is 600 nM for 1:1 mixture with 2.5 ng tRNA synthetase/Anap-CUA plasmid. As the volume of the oocyte nucleus is ∼40 nL, and it can tolerate 15-20 nL of injected volume (Lin-Moshier & Marchant, 2013), the final concentration of SIK inhibitor inside the oocyte nucleus was ∼150 nM.

First of all, *Ci-*VSP F401Anap was used to confirm reproducible effects in the initial optimization experiments. The most effective concentration of SIK inhibitor was determined to be 300 nM. Next, 300 nM of SIK inhibitor was co-injected to either the nucleus or cytoplasm of the oocytes, which were then incubated for different periods of time. This resulted in three test groups: (1) nuclear injection with 2 days incubation; (2) nuclear injection with 3 days incubation; and (3) cytoplasmic injection with 2 days incubation. Cytoplasmic injection needs concentration adjustment, since the volume of an oocyte is ∼1 μL. To make the concentration inside the oocyte 150 nM, the injected concentration was 3 μM. Control groups consisted of non-treated oocytes, incubated for either 2 or 3 days.

A follow-up confirmation experiment was done using the P2X2 A337Anap/R313W mutant, after the optimum concentration, injection method, and incubation days were determined from the *Ci*-VSP experiment. 300 nM of SIK inhibitor was co-injected into the nucleus of the oocyte. Oocytes were then incubated for 2-3 days after subsequent cytoplasmic co-injection of channel cRNA and Anap.

### Voltage-clamp fluorometry (VCF) recording

Oocytes for VCF-fUAA recording needed to be shielded from light exposure. Oocytes were placed in a recording chamber with the animal pole facing upward. For ATP-evoked current recording, a gap-free protocol was applied, with the holding potential at −80 mV. ATP was applied by perfusion system as described above. For voltage-evoked current recording, oocytes were held at +20 mV or at −40 mV in some cases. The step pulses were applied from +40 mV to −140 mV, +40 mV to −160 mV, or +80 mV to – 160 mV.

Two recordings (ATP application and voltage application) were performed separately in different oocytes. Meanwhile, VCF recordings in the absence and presence of ATP using voltage step pulses, for some mutants (A337Anap, R313F/A337Anap, R313W/A337Anap, and K308R/A337Anap), were performed in the same oocytes.

For voltage step application, ATP was applied directly. As bath volume was measured to be 600 μL, 20 μL ATP of 30 times higher concentration was applied directly to the bath solution. For *Ci*-VSP voltage-clamp recording, cells were clamped at −60 mV and the step pulses were applied from −80 mV to +160 mV every 3 seconds.

The fluorometric recordings were performed with an upright fluorescence microscope (Olympus BX51WI) equipped with a water immersion objective lens (Olympus XLUMPLAN FL 20x/1.00) to collect the emission light from the voltage-clamped oocytes. The light from a xenon arc lamp (L2194-01, Hamamatsu Photonics) was applied through a band-pass excitation filter (330-360 nm for Anap). In the case of the excitation of Anap to minimize photobleaching during ATP-application recording, the intensity of the excitation light was decreased to 1.5% by ND filters (U-25ND6 and U-25ND25 Olympus), whereas, for step-pulses recording, the intensity of the excitation light was decreased to 6% (U-25ND6 Olympus). Emitted light was passed through band pass emission filters (Brightline, Semrock) of 420–460 nm and 460–510 nm (Lee et al., 2009; Sakata et al., 2016). The emission signals were detected by two photomultipliers (H10722-110; Hamamatsu Photonics). The detected emission intensities were acquired by a Digidata 1332 (Axon Instruments) and Clampex 10.3 software (Molecular Devices) at 10 kHz for ATP application and 20 kHz for voltage application. In the case of *Ci*-VSP, the detected emission was acquired at 10 kHz. To improve the signal-to-noise ratio, VCF recording during step-pulse protocols was repeated 20 times for each sample for P2X2 in the presence of ATP, 5 times in the absence of ATP, and 3 times for *Ci*-VSP. Averaged data were used for data presentation and analysis.

### Data analysis

Two electrode voltage-clamp data were analyzed using Clampfit 10.5 software (Molecular Devices) and Igor Pro 5.01 (Wavemetrics). Analyses of conductance-voltage (G-1. V) relationship of P2X2 were obtained from tail current recordings at −60 mV and fitted to a two-state Boltzmann equation using Clampfit:

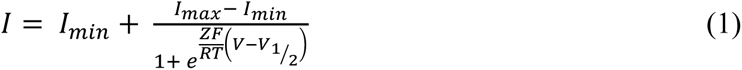

where I_min_ and I_max_ are defined as the limits of the amplitudes in fittings, Z is defined as the effective charge, V_1/2_ is the voltage of half activation, F is Faraday’s constant, and T is temperature in Kelvin.

In the case of P2X2, Normalized conductance-voltage (G-V) relationships were plotted using:

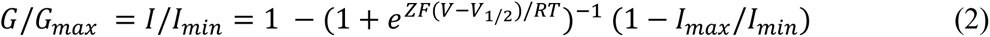

In the case of voltage-clamp fluorometry data, the gradual decline of fluorescence recording traces due to photobleaching was compensated by subtracting the expected time-lapse decrease in bleached component calculated from the trace’s bleaching rate (R) by assuming that the fluorescence is linear. Arithmetic operations were performed by Igor Pro 5.01 for ATP-evoked fluorescent signals.

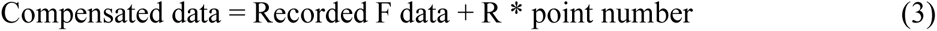

In the case of fluorescence traces from voltage application for both P2X2 and *Ci*-VSP, arithmetic operations were performed by Clampfit.

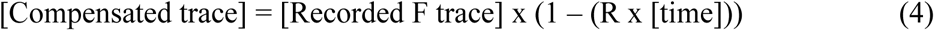

Where [time] is the value of the point given by Clampfit. All the compensated traces were then normalized by setting each baseline (F signal at −40 mV or at +20 mV depending on the holding potential) level to be 1 to calculate the % F change (ΔF/F; ΔF = F_−160mV_ - F_baseline_; F = F_baseline_). The fraction of ΔF_Slow_/F was calculated from the equation:

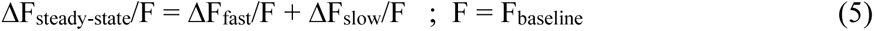

The data were expressed as mean±s.e.m with n indicating the number of samples.

### Statistical Analysis

Statistical analysis was performed by either one-way ANOVA, two-sample t-test, or paired t-test. Following one-way ANOVA, Tukey’s post-hoc test was applied. The data were expressed as mean ± s.e.m with n indicating the number of samples. Values p<0.05 were defined as statistically significant. *, **, *** denote values of p < 0.05, 0.01 and 0.001, respectively. All the statistical analysis and the bar graphs were performed and generated with OriginPro (OriginLab).

### Three-dimensional structural modelling of rat P2X2

Homology modelling was performed using a web-based environment for protein structure homology modelling SWISS-MODEL (Konstantin et al., 2006; Biasini et al., 2014) based upon sequence alignment of amino acids of *r*P2X2 (NM_053656) and the crystal structure of *h*P2X3 (Protein Data Bank accession number 5SVJ and 5SVK for closed and ATP-bound open state, respectively) (Mansoor et al., 2016). All the structural data presented in this study were generated using PyMOL molecular graphics system ver. 2.3.0 (Schrodinger LLC). Protein visualization was generated using Protter (Omasits et al., 2014).

## ACKNOWLEDGEMENTS

The authors thank Dr. Sakata and Prof. Okamura Y (Osaka University, Graduate School of Medicine) for the guidance of VCF experiments, all members in Kubo Laboratory for discussion, Ms. Naito C for technical support, and Dr. Collins A (Saba University, School of Medicine, Dutch Caribbean) for editing the manuscript.

---

**Figure 2—figure supplement 1.**
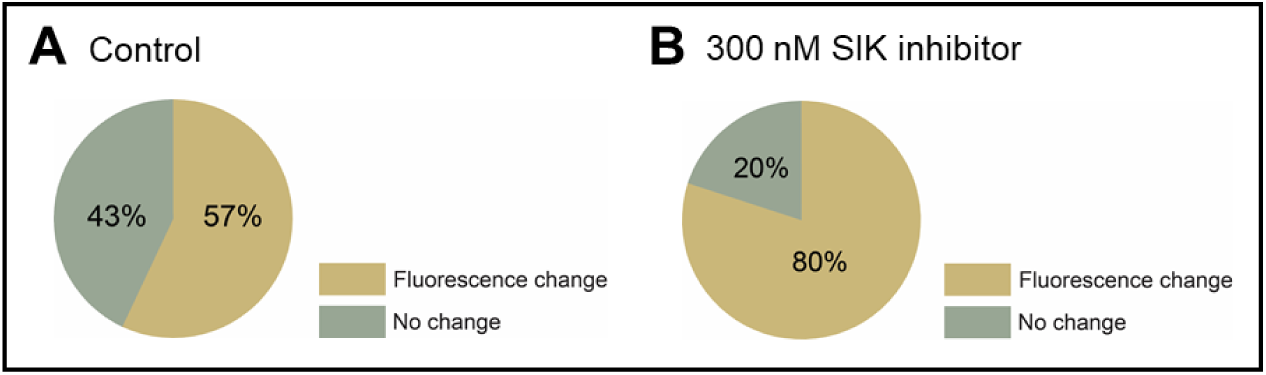
Effect of 300 nM SIK inhibitor application on the incidence of detectable Anap fluorescence signal change of P2X2 receptor. (**A, B**) Incidence of detectable changes of Anap fluorescence for control group (57%, n=7) and 300 nM SIK inhibitor application (80%, n=12), respectively. (**Source Data 1 Figure 2**)

**Figure 4—figure supplement 1.**
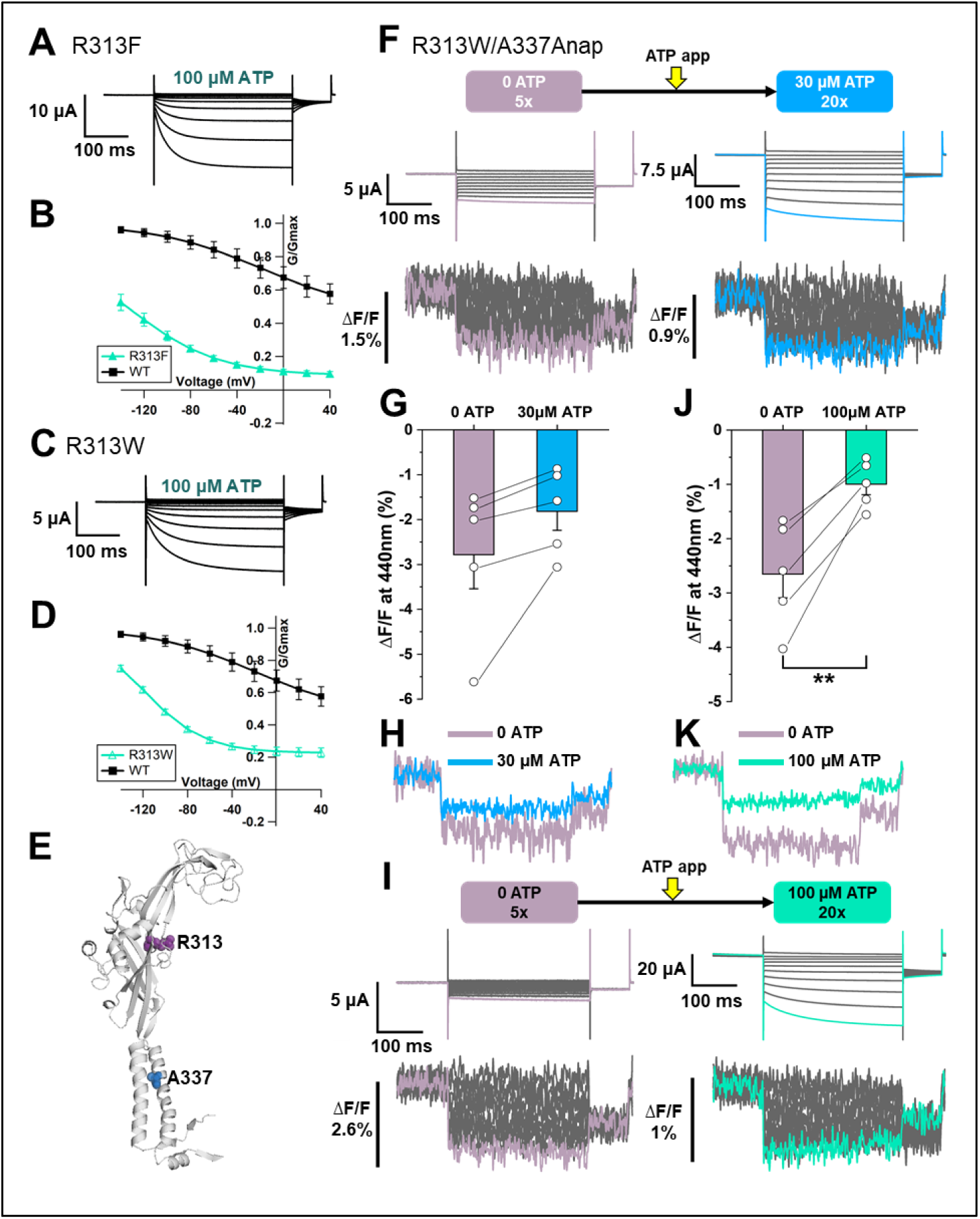
Voltage-clamp fluorometry of Anap-labeled P2X2 at A337 in TM2 evoked by hyperpolarization in the absence and presence of ATP. (**A**) Representative current traces of R313F upon application of 100 μM ATP (**B**) Comparison of G-V relationships between R313F (turquoise filled triangle) and wildtype (black filled square) in 100 μM ATP (n=3). (**C**) Representative current traces of R313W upon the application of 100 μM ATP. (**D**) G-V relationship comparison between R313W (turquoise open triangle) and wildtype (black filled square) in 100 μM ATP (n=3). Normalization was done based on the maximum conductance at the highest [ATP] (300 μM) for each construct. (**E**) Side view structural representation of the location of A337 (blue) and R313 (purple) in the ATP-bound open state (**F**) Representative current traces and fluorescence signal of VCF recordings of A337Anap/R313W in the absence of ATP (ΔF/F= 2.8%±0.7 at 440 nm, n=5) and 30 μM ATP (ΔF/F= 1.8%±0.4 at 440 nm, n=5). (**G**) Comparison of the fluorescence changes in the absence and in the presence of 30 μM ATP (p=0.072, paired t-test, n=5). (**H**) Superimposed fluorescence traces at −140 mV, in 0 ATP (light purple) and 30 μM ATP (blue). (**I**) Representative current traces and fluorescence signal of VCF recordings of A337/R313W, in the absence of ATP (ΔF/F= 2.6%±0.4 at 440 nm, n=5) and in 100 μM ATP (ΔF/F= 0.9%±0.2 at 440 nm, n=5) (**J**) Comparison of the fluorescence changes in the absence and in the presence of 100 μM ATP (** p≤0.01, p=0.0049, paired t-test, n=5). (**K**) Superimposed fluorescence traces at −140 mV, in 0 ATP (light purple) and 100 μM ATP (turquoise). All error bars are ± s.e.m centered on the mean. (**Source Data 1 Figure 4—figure supplement 1**)

**Figure 7—figure supplement 1.**
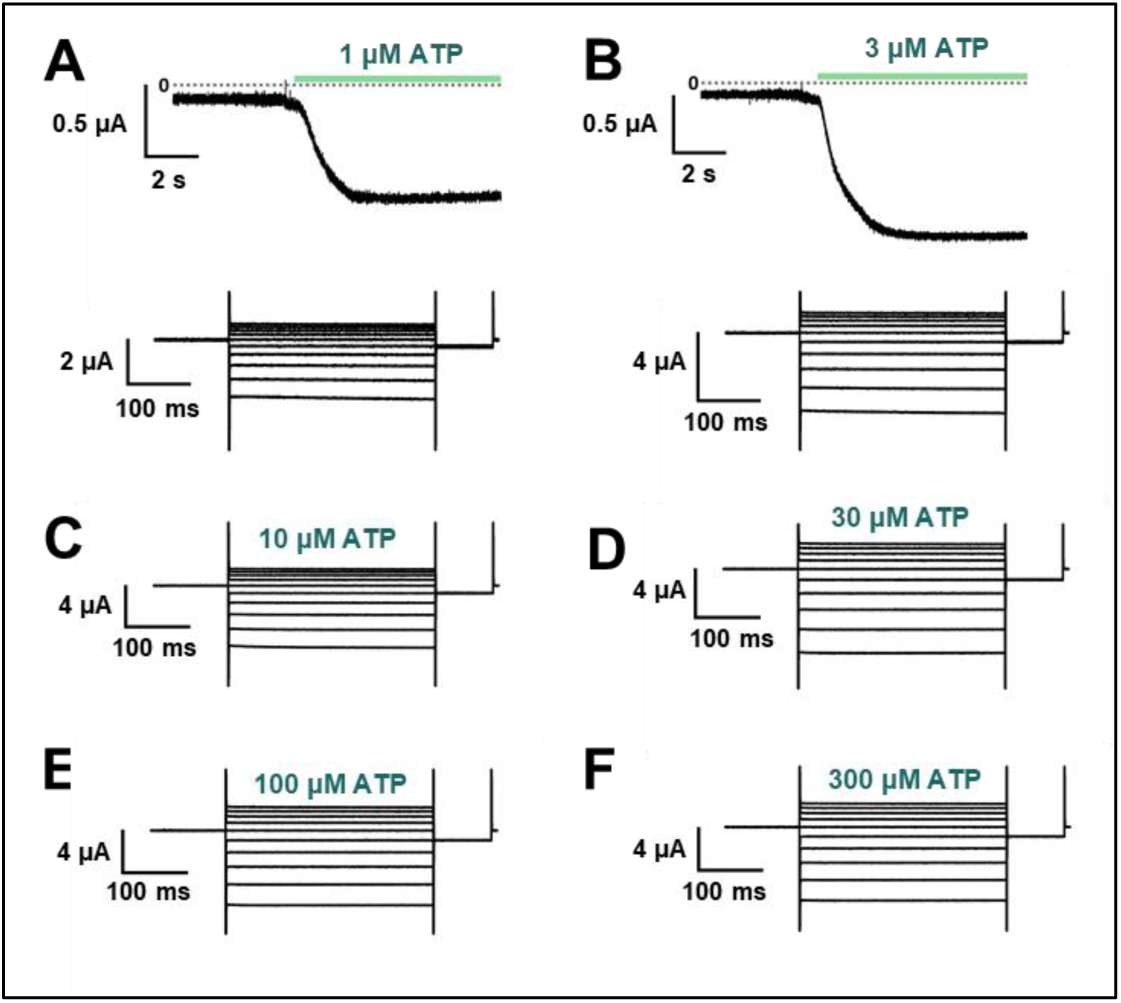
The effect of swapped mutation F44A/A337F. (**A – F**) Representative current traces of F44A/A337F upon various [ATP] application (1, 3, 10, 30, 100, 300 μM), followed by voltage application at each concentration (n=3). Voltage-dependent gating was almost absent, similarly to F44A (**Fig. 7E**).

**Supplementary Table 1.**
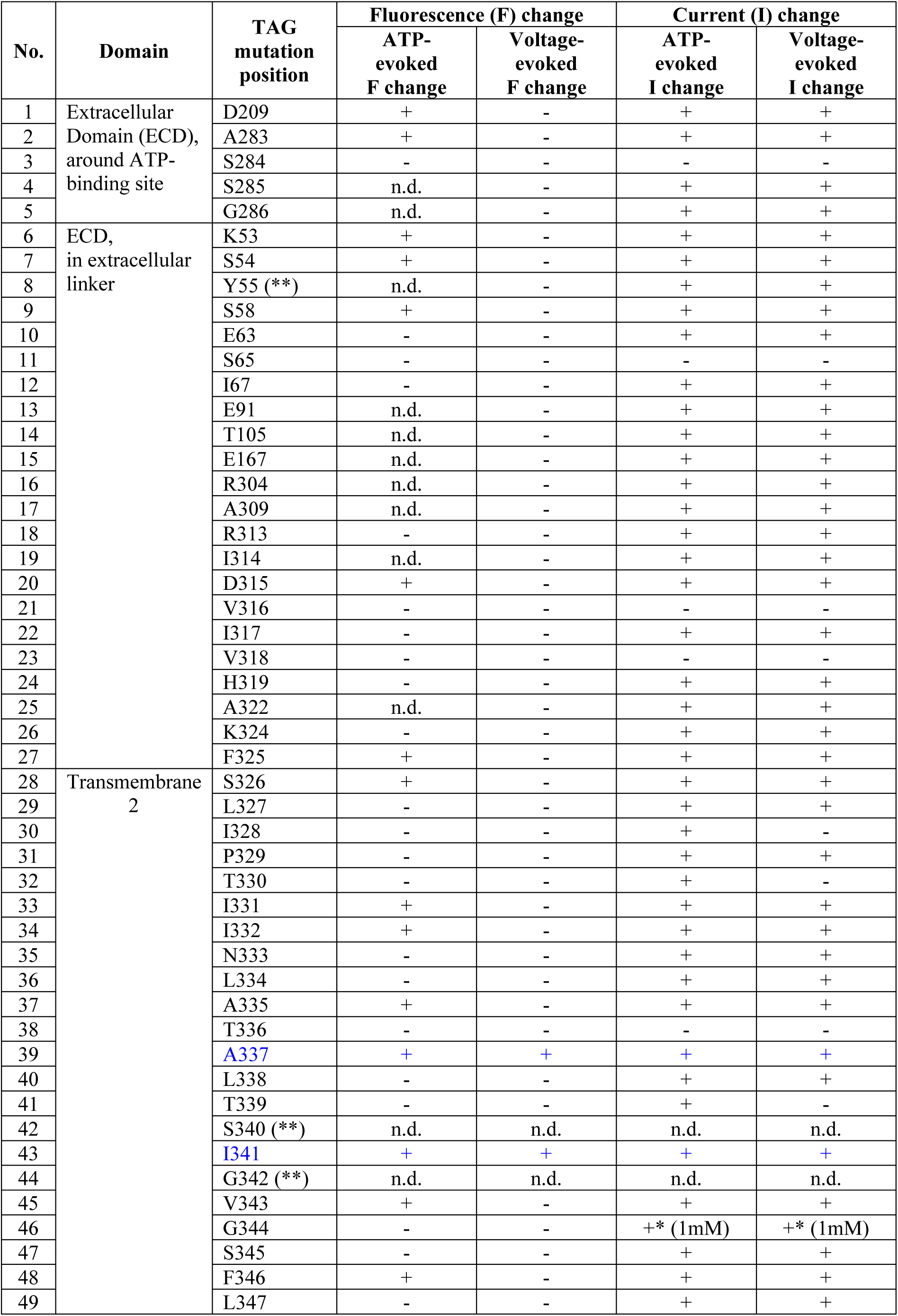
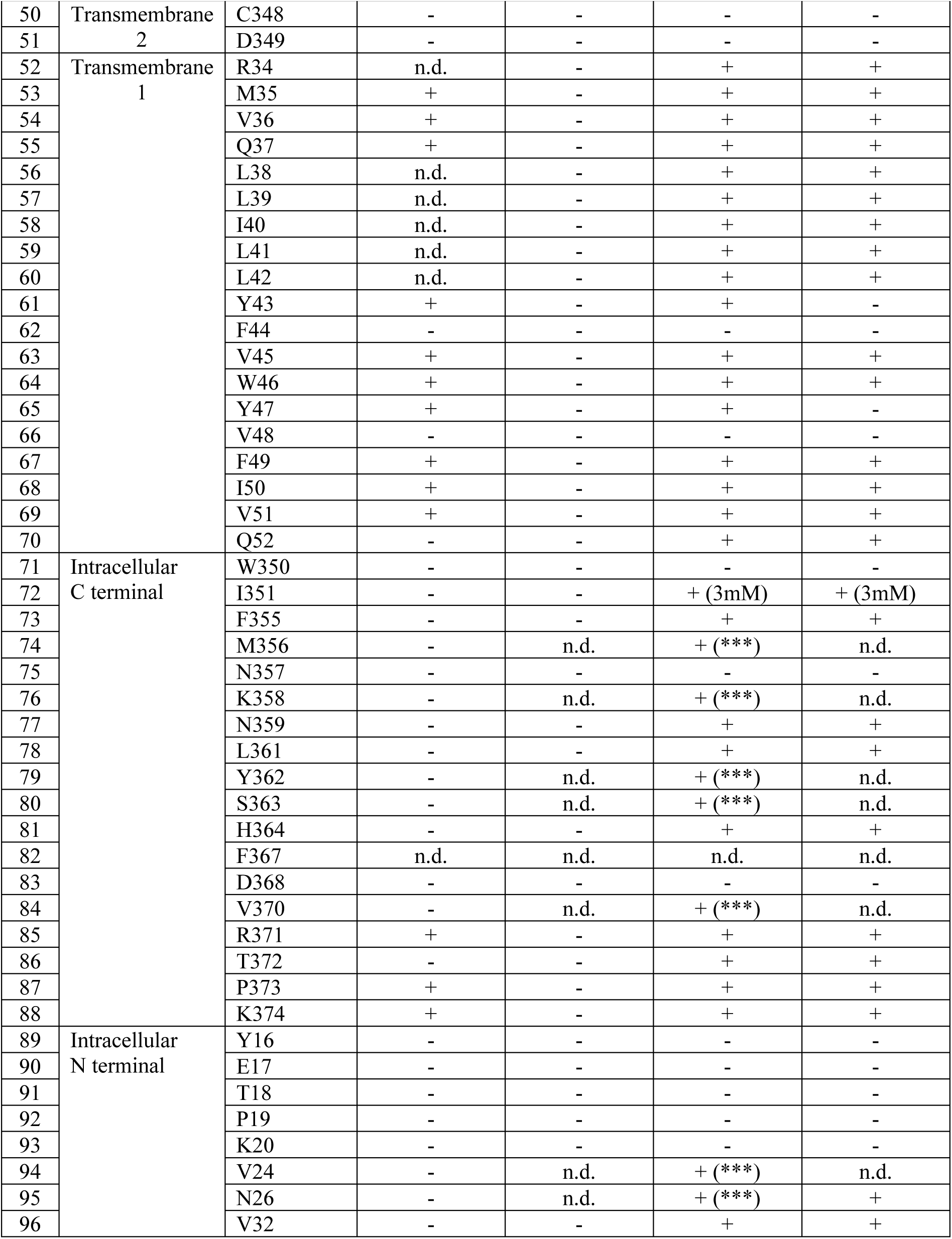
List of introduced TAG mutations in P2X2 receptor for VCF analyses. Mutations were introduced one at a time in 96 positions within the extracellular domain (ECD) near the ATP-binding site and extracellular linker, transmembrane domains (TMs), intracellular N-terminal, and intracellular C-terminal. ATP application ranging from 10 μM, 30 μM, or 100 μM unless otherwise stated. (+) indicates there was either ATP-evoked fluorescence (F) signal change, voltage-evoked F change, ATP-evoked current (I) change, or voltage-evoked I change. (-) indicates negative result. (**) indicates mutants which have a very low expression level so that the VCF analyses could not be performed. (***) indicates fast current decay. n.d. indicates not determined.

| No. | Domain | TAG mutation position | Fluorescence (F) change |  | Current (I) change |  |
| --- | --- | --- | --- | --- | --- | --- |
|  |  |  | ATP-evoked F change | Voltage-evoked F change | ATP-evoked I change | Voltage-evoked I change |
| 1 | Extracellular Domain (ECD), around ATP-binding site | D209 | + | - | + | + |
| 2 |  | A283 | + | - | + | + |
| 3 |  | S284 | - | - | - | - |
| 4 |  | S285 | n.d. | - | + | + |
| 5 |  | G286 | n.d. | - | + | + |
| 6 | ECD, in extracellular linker | K53 | + | - | + | + |
| 7 |  | S54 | + | - | + | + |
| 8 |  | Y55 (**) | n.d. | - | + | + |
| 9 |  | S58 | + | - | + | + |
| 10 |  | E63 | - | - | + | + |
| 11 |  | S65 | - | - | - | - |
| 12 |  | I67 | - | - | + | + |
| 13 |  | E91 | n.d. | - | + | + |
| 14 |  | T105 | n.d. | - | + | + |
| 15 |  | E167 | n.d. | - | + | + |
| 16 |  | R304 | n.d. | - | + | + |
| 17 |  | A309 | n.d. | - | + | + |
| 18 |  | R313 | - | - | + | + |
| 19 |  | I314 | n.d. | - | + | + |
| 20 |  | D315 | + | - | + | + |
| 21 |  | V316 | - | - | - | - |
| 22 |  | I317 | - | - | + | + |
| 23 |  | V318 | - | - | - | - |
| 24 |  | H319 | - | - | + | + |
| 25 |  | A322 | n.d. | - | + | + |
| 26 |  | K324 | - | - | + | + |
| 27 |  | F325 | + | - | + | + |
| 28 | Transmembrane 2 | S326 | + | - | + | + |
| 29 |  | L327 | - | - | + | + |
| 30 |  | I328 | - | - | + | - |
| 31 |  | P329 | - | - | + | + |
| 32 |  | T330 | - | - | + | - |
| 33 |  | I331 | + | - | + | + |
| 34 |  | I332 | + | - | + | + |
| 35 |  | N333 | - | - | + | + |
| 36 |  | L334 | - | - | + | + |
| 37 |  | A335 | + | - | + | + |
| 38 |  | T336 | - | - | - | - |
| 39 |  | A337 | + | + | + | + |
| 40 |  | L338 | - | - | + | + |
| 41 |  | T339 | - | - | + | - |
| 42 |  | S340 (**) | n.d. | n.d. | n.d. | n.d. |
| 43 |  | I341 | + | + | + | + |
| 44 |  | G342 (**) | n.d. | n.d. | n.d. | n.d. |
| 45 |  | V343 | + | - | + | + |
| 46 |  | G344 | - | - | +* (1mM) | +* (1mM) |
| 47 |  | S345 | - | - | + | + |
| 48 |  | F346 | + | - | + | + |
| 49 |  | L347 | - | - | + | + |
|  |  |  | ATP-evoked F change | Voltage-evoked F change | ATP-evoked I change | Voltage-evoked I change |
| 50 | Transmembrane 2 | C348 | - | - | - | - |
| 51 |  | D349 | - | - | - | - |
| 52 | Transmembrane 1 | R34 | n.d. | - | + | + |
| 53 |  | M35 | + | - | + | + |
| 54 |  | V36 | + | - | + | + |
| 55 |  | Q37 | + | - | + | + |
| 56 |  | L38 | n.d. | - | + | + |
| 57 |  | L39 | n.d. | - | + | + |
| 58 |  | I40 | n.d. | - | + | + |
| 59 |  | L41 | n.d. | - | + | + |
| 60 |  | L42 | n.d. | - | + | + |
| 61 |  | Y43 | + | - | + | - |
| 62 |  | F44 | - | - | - | - |
| 63 |  | V45 | + | - | + | + |
| 64 |  | W46 | + | - | + | + |
| 65 |  | Y47 | + | - | + | - |
| 66 |  | V48 | - | - | - | - |
| 67 |  | F49 | + | - | + | + |
| 68 |  | I50 | + | - | + | + |
| 69 |  | V51 | + | - | + | + |
| 70 |  | Q52 | - | - | + | + |
| 71 | Intracellular C terminal | W350 | - | - | - | - |
| 72 |  | I351 | - | - | + (3mM) | + (3mM) |
| 73 |  | F355 | - | - | + | + |
| 74 |  | M356 | - | n.d. | + (***) | n.d. |
| 75 |  | N357 | - | - | - | - |
| 76 |  | K358 | - | n.d. | + (***) | n.d. |
| 77 |  | N359 | - | - | + | + |
| 78 |  | L361 | - | - | + | + |
| 79 |  | Y362 | - | n.d. | + (***) | n.d. |
| 80 |  | S363 | - | n.d. | + (***) | n.d. |
| 81 |  | H364 | - | - | + | + |
| 82 |  | F367 | n.d. | n.d. | n.d. | n.d. |
| 83 |  | D368 | - | - | - | - |
| 84 |  | V370 | - | n.d. | + (***) | n.d. |
| 85 |  | R371 | + | - | + | + |
| 86 |  | T372 | - | - | + | + |
| 87 |  | P373 | + | - | + | + |
| 88 |  | K374 | + | - | + | + |
| 89 | Intracellular N terminal | Y16 | - | - | - | - |
| 90 |  | E17 | - | - | - | - |
| 91 |  | T18 | - | - | - | - |
| 92 |  | P19 | - | - | - | - |
| 93 |  | K20 | - | - | - | - |
| 94 |  | V24 | - | n.d. | + (***) | n.d. |
| 95 |  | N26 | - | n.d. | + (***) | + |
| 96 |  | V32 | - | - | + | + |

## REFERENCES

Asamoah OK, Wuskell JP, Loew LM, Bezanilla F. 2003. A Fluorometric Approach to Local Electric Field Measurements in a Voltage-Gated Ion Channel. Neuron, 37(1), 85–98. doi:https://doi.org/10.1016/S0896-6273(02)01126-1

Barchad-Avitzur O, Priest MF, Dekel N, Bezanilla F, Parnas H, Ben-Chaim Y. 2016. A Novel Voltage Sensor in the Orthosteric Binding Site of the M2 Muscarinic Receptor. Biophysical Journal, 111(7), 1396–1408. doi:10.1016/j.bpj.2016.08.035

Ben-Chaim Y, Chanda B, Dascal N, Bezanilla F, Parnas I, Parnas H. 2006. Movement of ‘gating charge’ is coupled to ligand binding in a G-protein-coupled receptor. Nature, 444(7115), 106–109. doi:10.1038/nature05259

Bezanilla F. 2008. How membrane proteins sense voltage. Nature Reviews Molecular Cell Biology, 9, 323. doi:10.1038/nrm2376

Biasini M, Bienert S, Waterhouse A, Arnold K, Studer G, Schmidt T, Bordoli L, Schwede T. 2014. SWISS MODEL: modelling protein tertiary and quaternary structure using evolutionary information. Nucleic Acid Research, 42, 252–258. doi:10.1093/nar/gku340

Brake AJ, Wagenbach MJ, Julius D. 1994. New structural motif for ligand-gated ion channels defined by an ionotropic ATP receptor. Nature, 371(6497), 519–523. doi:10.1038/371519a0

Bublitz GU, Boxer SG. 1997. STARK SPECTROSCOPY: Applications in Chemistry, Biology, and Materials Science. Annual Review of Physical Chemistry, 48(1), 213–242. doi:10.1146/annurev.physchem.48.1.213

Burnstock G. (2003). Introduction: ATP and Its Metabolites as Potent Extracellular Agents Current Topics in Membranes (Vol. 54, pp. 1–27): Academic Press.

Cao L, Broomhead HE, Young MT, North RA. 2009. Polar residues in the second transmembrane domain of the rat P2X2 receptor that affect spontaneous gating, unitary conductance, and rectification. J Neurosci, 29(45), 14257–14264. doi:10.1523/JNEUROSCI.4403-09.2009

Cao L, Young MT, Broomhead HE, Fountain SJ, North RA. 2007. Thr339-to-serine substitution in rat P2X2 receptor second transmembrane domain causes constitutive opening and indicates a gating role for Lys308. J Neurosci, 27(47), 12916–12923. doi:10.1523/JNEUROSCI.4036-07.2007

Cha A, Bezanilla F. 1997. Characterizing Voltage-Dependent Conformational Changes in the ShakerK+ Channel with Fluorescence. Neuron, 19(5), 1127–1140. doi:https://doi.org/10.1016/S0896-6273(00)80403-1

Chatterjee A, Guo J, Lee HS, Schultz PG. 2013. A genetically encoded fluorescent probe in mammalian cells. J Am Chem Soc, 135(34), 12540–12543. doi:10.1021/ja4059553

Dekel N, Priest MF, Parnas H, Parnas I, Bezanilla F. 2012. Depolarization induces a conformational change in the binding site region of the M2 muscarinic receptor. Proc Natl Acad Sci U S A, 109(1), 285–290. doi:10.1073/pnas.1119424109

Ding S, Sachs F. 2000. Inactivation of P2X2 purinoceptors by divalent cations. J Physiol, 522 *Pt 2*(Pt 2), 199–214. doi:10.1111/j.1469-7793.2000.t01-1-00199.x

Ennion S, Hagan S, Evans RJ. 2000. The role of positively charged amino acids in ATP recognition by human P2X(1) receptors. J Biol Chem, 275(38), 29361–29367. doi:10.1074/jbc.M003637200

Fujiwara Y, Keceli B, Nakajo K, Kubo Y. 2009. Voltage- and [ATP]-dependent gating of the P2X(2) ATP receptor channel. J Gen Physiol, 133(1), 93–109. doi:10.1085/jgp.200810002

Fujiwara Y, Kubo Y. 2004. Density-dependent changes of the pore properties of the P2X2 receptor channel. J Physiol, 558(Pt 1), 31–43. doi:10.1113/jphysiol.2004.064568

Habermacher C, Martz A, Calimet N, Lemoine D, Peverini L, Specht A, Cecchini M, Grutter T. 2016. Photo-switchable tweezers illuminate pore-opening motions of an ATP-gated P2X ion channel. Elife, 5, e11050. doi:10.7554/eLife.11050

Haines WR, Migita K, Cox JA, Egan TM, Voigt MM. 2001. The first transmembrane domain of the P2X receptor subunit participates in the agonist-induced gating of the channel. J Biol Chem, 276(35), 32793–32798. doi:10.1074/jbc.M104216200

Hattori M, Gouaux E. 2012. Molecular mechanism of ATP binding and ion channel activation in P2X receptors. Nature, 485(7397), 207–212. doi:10.1038/nature11010

Heymann G, Dai J, Li M, Silberberg SD, Zhou HX, Swartz KJ. 2013. Inter- and intrasubunit interactions between transmembrane helices in the open state of P2X receptor channels. Proc Natl Acad Sci U S A, 110(42), E4045–4054. doi:10.1073/pnas.1311071110

Housley GD, Morton-Jones R, Vlajkovic SM, Telang RS, Paramananthasivam V, Tadros SF, Wong ACY, Froud KE, Cederholm JME, Sivakumaran Y, Snguanwongchai P, Khakh BS, Cockayne DA, Thorne PR, Ryan AF. 2013. ATP-gated ion channels mediate adaptation to elevated sound levels. Proc Natl Acad Sci U S A, 110(18), 7494–7499. doi:10.1073/pnas.1222295110

Jiang L-H, Kim M, Spelta V, Bo X, Surprenant A, North RA. 2003. Subunit Arrangement in P2X Receptors. The Journal of Neuroscience, 23(26), 8903. doi:10.1523/JNEUROSCI.23-26-08903.2003

Jiang LH, Rassendren F, Spelta V, Surprenant A, North RA. 2001. Amino acid residues involved in gating identified in the first membrane-spanning domain of the rat P2X(2) receptor. J Biol Chem, 276(18), 14902–14908. doi:10.1074/jbc.M011327200

Jiang LH, Rassendren F, Surprenant A, North RA. 2000. Identification of amino acid residues contributing to the ATP-binding site of a purinergic P2X receptor. J Biol Chem, 275(44), 34190–34196. doi:10.1074/jbc.M005481200

Jiang Y, Lee A, Chen J, Ruta V, Cadene M, Chait BT, MacKinnon R. 2003. X-ray structure of a voltage-dependent K+ channel. Nature, 423(6935), 33–41. doi:10.1038/nature01580

Kalstrup T, Blunck R. 2013. Dynamics of internal pore opening in K_V_ channels probed by a fluorescent unnatural amino acid. Proc Natl Acad Sci U S A, 110(20), 8272–8277. doi:10.1073/pnas.1220398110

Kalstrup T, Blunck R. 2018. S4-S5 linker movement during activation and inactivation in voltage-gated K(+) channels. Proc Natl Acad Sci U S A, 115(29), E6751–E6759. doi:10.1073/pnas.1719105115

Kawate T, Michel JC, Birdsong WT, Gouaux E. 2009. Crystal structure of the ATP-gated P2X(4) ion channel in the closed state. Nature, 460(7255), 592–598. doi:10.1038/nature08198

Keceli B, Kubo Y. 2009. Functional and structural identification of amino acid residues of the P2X2 receptor channel critical for the voltage- and [ATP]-dependent gating. J Physiol, 587(Pt 24), 5801–5818. doi:10.1113/jphysiol.2009.182824

Keceli B, Kubo Y. 2014. Signal transmission within the P2X2 trimeric receptor. J Gen Physiol, 143(6), 761–782. doi:10.1085/jgp.201411166

Khakh BS, Egan TM. 2005. Contribution of transmembrane regions to ATP-gated P2X2 channel permeability dynamics. J Biol Chem, 280(7), 6118–6129. doi:10.1074/jbc.M411324200

Klippenstein V, Mony L, Paoletti P. 2018. Probing Ion Channel Structure and Function Using Light-Sensitive Amino Acids. Trends Biochem Sci, 43(6), 436–451. doi:10.1016/j.tibs.2018.02.012

Klymchenko AS, Demchenko AP. 2002. Electrochromic Modulation of Excited-State Intramolecular Proton Transfer: The New Principle in Design of Fluorescence Sensors. Journal of the American Chemical Society, 124(41), 12372–12379. doi:10.1021/ja027669l

Klymchenko AS, Stoeckel H, Takeda K, Mély Y. 2006. Fluorescent Probe Based on Intramolecular Proton Transfer for Fast Ratiometric Measurement of Cellular Transmembrane Potential. The Journal of Physical Chemistry B, 110(27), 13624–13632. doi:10.1021/jp062385z

Konstantin A, Lorenza B, Kopp J, Torsten S. 2006. The SWISS Model workspace: a web-based environment for protein structure homology modelling. Bioinformatics, 22, 195–201.

Kracun S, Chaptal V, Abramson J, Khakh BS. 2010. Gated access to the pore of a P2X receptor: structural implications for closed-open transitions. J Biol Chem, 285(13), 10110–10121. doi:10.1074/jbc.M109.089185

Lee EEL, Bezanilla F. 2019. Methodological improvements for fluorescence recordings in Xenopus laevis oocytes. J Gen Physiol, 151(2), 264–272. doi:10.1085/jgp.201812189

Lee HS, Guo J, Lemke EA, Dimla RD, Schultz PG. 2009. Genetic incorporation of a small, environmentally sensitive, fluorescent probe into proteins in Saccharomyces cerevisiae. J Am Chem Soc, 131(36), 12921–12923. doi:10.1021/ja904896s

Li M, Chang TH, Silberberg SD, Swartz KJ. 2008. Gating the pore of P2X receptor channels. Nat Neurosci, 11(8), 883–887. doi:10.1038/nn.2151

Li M, Kawate T, Silberberg SD, Swartz KJ. 2010. Pore-opening mechanism in trimeric P2X receptor channels. Nat Commun, 1, 44. doi:10.1038/ncomms1048

Li Z, Migita K, Samways DSK, Voigt MM, Egan TM. 2004. Gain and Loss of Channel Function by Alanine Substitutions in the Transmembrane Segments of the Rat ATP-Gated P2X2 Receptor. The Journal of Neuroscience, 24(33), 7378–7386. doi:10.1523/jneurosci.1423-04.2004

Lin-Moshier Y, Marchant JS. 2013. Nuclear microinjection to assess how heterologously expressed proteins impact Ca2+ signals in Xenopus oocytes. Cold Spring Harb Protoc, 2013(3). doi:10.1101/pdb.prot072785

Loew LM. 1982. Design and characterization of electrochromic membrane probes. Journal of Biochemical and Biophysical Methods, 6(3), 243–260. doi:https://doi.org/10.1016/0165-022X(82)90047-1

Mannuzzu LM, Moronne MM, Isacoff EY. 1996. Direct Physical Measure of Conformational Rearrangement Underlying Potassium Channel Gating. Science, 271(5246), 213–216. doi:10.1126/science.271.5246.213

Mansoor SE, Lu W, Oosterheert W, Shekhar M, Tajkhorshid E, Gouaux E. 2016. X-ray structures define human P2X(3) receptor gating cycle and antagonist action. Nature, 538(7623), 66–71. doi:10.1038/nature19367

McCarthy AE, Yoshioka C, Mansoor SE. 2019. Full-Length P2X7 Structures Reveal How Palmitoylation Prevents Channel Desensitization. Cell, 179(3), 659–670.e613. doi:https://doi.org/10.1016/j.cell.2019.09.017

Mujahid N, Liang Y, Murakami R, Choi HG, Dobry AS, Wang J, Suita Y, Weng QY, Allouche J, Kemeny LV, Hermann AL, Roider EM, Gray NS, Fisher DE. 2017. A UV-Independent Topical Small-Molecule Approach for Melanin Production in Human Skin. Cell Reports, 19(11), 2177–2184. doi:https://doi.org/10.1016/j.celrep.2017.05.042

Nakajo K, Kubo Y. 2014. Steric hindrance between S4 and S5 of the KCNQ1/KCNE1 channel hampers pore opening. Nat Commun, 5, 4100. doi:10.1038/ncomms5100

Nakazawa K, Inoue K, Ohno Y. 1998. An asparagine residue regulating conductance through P2X2 receptor/channels. European Journal of Pharmacology, 347(1), 141–144. doi:https://doi.org/10.1016/S0014-2999(98)00207-6

Nakazawa K, Liu M, Inoue K, Ohno Y. 1997. Voltage-Dependent Gating of ATP-Activated Channels in PC12 Cells. Journal of Neurophysiology, 78(2), 884–890. doi:10.1152/jn.1997.78.2.884

Nakazawa K, Ohno Y. 2005. Characterization of voltage-dependent gating of P2X2 receptor/channel. European Journal of Pharmacology, 508(1), 23–30. doi:https://doi.org/10.1016/j.ejphar.2004.12.005

Navarro-Polanco RA, Moreno Galindo EG, Ferrer-Villada T, Arias M, Rigby JR, Sanchez-Chapula JA, Tristani-Firouzi M. 2011. Conformational changes in the M2 muscarinic receptor induced by membrane voltage and agonist binding. J Physiol, 589(Pt 7), 1741–1753. doi:10.1113/jphysiol.2010.204107

North RA. 2002. Molecular Physiology of P2X Receptors. Physiological Reviews, 82(4), 1013–1067. doi:10.1152/physrev.00015.2002

Omasits U, Ahrens CH, Müller S, Wollscheid B. 2014. Protter: interactive protein feature visualization and integration with experimental proteomic data. Bioinformatics(15 March 2014), 884–886. doi:10.1093/bioinformatics/btt607.

Papp F, Lomash S, Szilagyi O, Babikow E, Smith J, Chang TH, Bahamonde MI, Toombes GES, Swartz KJ. 2019. TMEM266 is a functional voltage sensor regulated by extracellular Zn(2). Elife, 8. doi:10.7554/eLife.42372

Pless SA, Lynch JW. 2008. Illuminating the Structure and Function of Cys-Loop Receptors. Clinical and Experimental Pharmacology and Physiology, 35(10), 1137–1142. doi:10.1111/j.1440-1681.2008.04954.x

Radford KM, Virginio C, Surprenant A, North RA, Kawashima E. 1997. Baculovirus Expression Provides Direct Evidence for Heteromeric Assembly of P2X2 and P2X3 Receptors. The Journal of Neuroscience, 17(17), 6529. doi:10.1523/JNEUROSCI.17-17-06529.1997

Roberts JA, Vial C, Digby HR, Agboh KC, Wen H, Atterbury-Thomas A, Evans RJ. 2006. Molecular properties of P2X receptors. Pflugers Arch, 452(5), 486–500. doi:10.1007/s00424-006-0073-6

Sakata S, Jinno Y, Kawanabe A, Okamura Y. 2016. Voltage-dependent motion of the catalytic region of voltage-sensing phosphatase monitored by a fluorescent amino acid. Proc Natl Acad Sci U S A, 113(27), 7521–7526. doi:10.1073/pnas.1604218113

Samways DS, Migita K, Li Z, Egan TM. 2008. On the role of the first transmembrane domain in cation permeability and flux of the ATP-gated P2X2 receptor. J Biol Chem, 283(8), 5110–5117. doi:10.1074/jbc.M708713200

Schewe M, Nematian-Ardestani E, Sun H, Musinszki M, Cordeiro S, Bucci G, de Groot BL, Tucker SJ, Rapedius M, Baukrowitz T. 2016. A Non-canonical Voltage-Sensing Mechanism Controls Gating in K2P K(+) Channels. Cell, 164(5), 937–949. doi:10.1016/j.cell.2016.02.002

Stelmashenko O, Compan V, Browne LE, North RA. 2014. Ectodomain movements of an ATP-gated ion channel (P2X2 receptor) probed by disulfide locking. J Biol Chem, 289(14), 9909–9917. doi:10.1074/jbc.M113.542811

Swartz KJ. 2008. Sensing voltage across lipid membranes. Nature, 456, 891. doi:10.1038/nature07620

Talwar S, Lynch JW. 2015. Investigating ion channel conformational changes using voltage clamp fluorometry. Neuropharmacology, 98, 3–12. doi:10.1016/j.neuropharm.2015.03.018

Valera S, Hussy N, Evans RJ, Adami N, North RA, Surprenant A, Buell G. 1994. A new class of ligand-gated ion channel defined by P2X receptor for extracellular ATP. Nature, 371(6497), 516–519. doi:10.1038/371516a0

Yan D, Zhu Y, Walsh T, Xie D, Yuan H, Sirmaci A, Fujikawa T, Wong ACY, Loh TL, Du L, Grati Mh, Vlajkovic SM, Blanton S, Ryan AF, Chen Z-Y, Thorne PR, Kachar B, Tekin M, Zhao H-B, Housley GD, King M-C, Liu XZ. 2013. Mutation of the ATP-gated P2X2 receptor leads to progressive hearing loss and increased susceptibility to noise. Proc Natl Acad Sci U S A, 110(6), 2228–2233. doi:10.1073/pnas.1222285110

Zhang J, Davidson RM, Wei M-d, Loew LM. 1998. Membrane Electric Properties by Combined Patch Clamp and Fluorescence Ratio Imaging in Single Neurons. Biophysical Journal, 74(1), 48–53. doi:https://doi.org/10.1016/S0006-3495(98)77765-3

Zhou Z, Hume RI. 1998. Two mechanisms for inward rectification of current flow through the purinoceptor P2X2 class of ATP-gated channels. J Physiol, 507 (Pt 2), 353–364. doi:10.1111/j.1469-7793.1998.353bt.x

